# NodoMap: a spatio-cellular map of the mouse nodose ganglia

**DOI:** 10.1101/2025.05.11.653249

**Authors:** Sijing Cheng, Georgina K. C. Dowsett, Kara Rainbow, Mariana Norton, Anna G. Roberts, Phyllis Phuah, Gavin A. Bewick, Brian Y.H. Lam, Giles S.H. Yeo, Kevin G. Murphy

## Abstract

The vagus nerve is a key component of the parasympathetic nervous system, innervating multiple abdominal organs to monitor and regulate their function. It forms the main neural pathway between the gastrointestinal tract and the brain, playing a major role in the regulation of energy homeostasis. The cell bodies for vagal sensory neurons reside in the nodose ganglia, with the left and right ganglia reported to have distinct roles in food intake and reward. Here, we have integrated our own single nucleus RNA sequencing data with multiple publicly available datasets to create a database of 108,482 nuclei and cells, and combined this with spatial transcriptomics to present a spatio-cellular transcriptional map of the mouse nodose ganglia, the ‘NodoMap’. Nodose ganglia neuronal cells clustered into twenty-two different subtypes, all found in both left and right nodose ganglia, but with significant differences in gene expression between left and right ganglia across multiple neuronal subtypes. Overnight fasting modulated gene expression across specific neuronal subtypes, including nutrient responsive pathways. Spatial transcriptomics showed that while vagal neuronal types were highly interspersed, patterns of organisation into cellular ‘neighbourhoods’ could be observed, with neighbourhoods identified of predominantly non-neuronal cells and of different neuronal populations accompanied by glial-like cells. Thus, NodoMap provides a detailed atlas of the mouse nodose ganglia in a spatial context, providing a platform for vagovagal neurocircuit analysis, and serving as an important resource to identify targets for pharmacotherapies for metabolic disease.

## Introduction

The vagus nerve is a major component of the parasympathetic nervous system and transmits information crucial to an array of physiological functions between the brain and peripheral organs. Vagal afferent neurons signal to the brainstem, where they can act on interneurons, which either modulate other circuits, or input into local vagal efferent neurons, forming vagovagal neurocircuits that detect and respond to sensory inputs to regulate motor functions^1,2^. The vagus is an integral part, and the main neuronal pathway, of the gut-brain axis, which regulates appetite and gut function. The subdiaphragmatic sensory afferent fibres of the vagus nerve innervate the gastrointestinal tract, where vagal mechanoreceptors and chemoreceptors can detect gastrointestinal distension, as well as the presence and absorption of nutrients, both directly, and via their effects on the release of gastrointestinal hormones^3,4^. Information regarding luminal pressure, nutrient, and hormonal signalling can thus be promptly transduced to the brain, and integrated to regulate appetite and gastrointestinal motility. Vagus nerve stimulation can also be used to treat specific neurological conditions^5–8^, and has been suggested to be potentially useful in other diseases, including obesity^9–12^.

The cell bodies of vagal afferent fibres reside in the nodose ganglia, located at the base of skull^13^. Nodose ganglia neurons are heterogeneous and can be separated into either mechanosensory or nociceptor types^14^. In the last decade, new technologies have revolutionised our understanding of the vagus^15–20^. The nodose ganglia form part of a nodose-jugular ganglia complex in mice^21^, and mouse models have been widely used for preclinical studies of vagal signalling. However, there is no unified RNA sequencing database at single cell or nucleus level to characterise vagal afferents, and the spatial organisation of these neurons is largely unexplored. While different signals have been shown to be transduced through the left or right nodose, and transcriptional changes in specific genes that occur in the nodose ganglia in response to alterations in nutritional state have been reported^22–27^, a comprehensive comparison of the neuronal populations on both sides has yet to be performed^18,28,29^. We therefore aimed to produce a genetic map of the nodose ganglia as a tool to help understand and further interrogate the role of specific vagal neuronal populations.

## RESULTS

Several single-cell RNA sequencing (scRNASeq) studies have been performed on murine nodose ganglia^14,15,19,28,30^. However, single-cell dissociation has to be performed on fresh tissue, is time consuming, risks cell-type selection bias, and can often induce transcriptional perturbation^31^. In contrast, single-nucleus RNA sequencing (snRNASeq) can be performed on frozen samples, and nuclear RNA has been found to be more stable than cytoplasmic RNA in brain samples^32^. Crucially, several studies have shown that the transcriptomic profile of murine brain tissues generated from snRNASeq is comparable to that generated by scRNASeq^31–34^. Thus, we performed snRNASeq on left and right nodose ganglia taken from mice fed *ad libitum* or fasted overnight. To establish a clear transcriptomic definition of the nodose ganglia, we integrated our in-house snRNASeq data with four other published datasets (Figure S1A)^14,15,28,30^, in order to create a unified reference atlas, ‘NodoMap’ (Figure 1A), comprising a total of 108,482 cells, and characterising corresponding cell types across different studies^35^.

**Figure 1:**
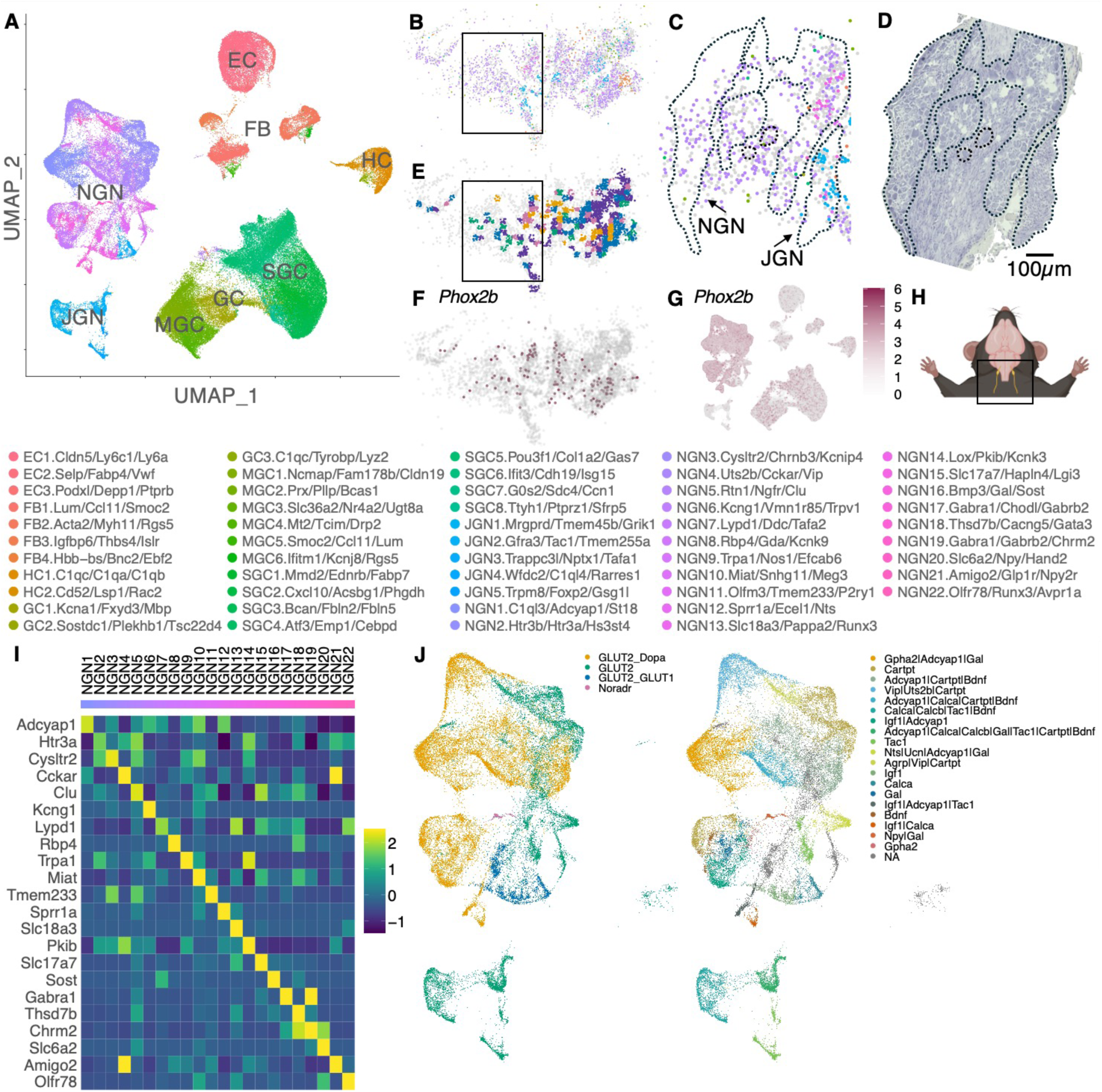
A spatio-cellular atlas of the mouse nodose ganglion. **(A)** A UMAP plot of the single cell/nuclei dataset, coloured by cluster and with cluster names written below. We integrated 4 publicly available and 1 in-house sc/snRNASeq datasets of the mouse nodose ganglia to create a database of 108,482 cells. Of this, 33,451 were neurons and 50,485 were ‘satellite glial cells’ and ‘myelinating glial cells’. There were also 10,709 endothelial cells, 8874 fibroblasts and 4963 hemopoietic cells. Clusters were annotated based on cell type and the top 3 marker genes with the highest specificity for that cluster. **(B)** spatial transcriptomics of the nodose ganglion. To identify the cell type present underneath each spot we integrated the sc/snRNASeq atlas together with the spatial transcriptomics data using RCTD (see methods), to predict the cell type and number of cells present at each bead. slide-seq data is presented for 1 tile of 3 left nodose ganglia sections; spots represent beads where RNA transcripts were detected. Spots are coloured based on the predicted cell type present at each bead. Beads were annotated with a cell type only if the RCTD pipeline identified them as singlets. Colours match the sc/snRNASeq cluster colours. **(C)** a zoomed-in image of the square section from panel B, and **(D)** Haematoxylin staining of an adjacent section of the same region. In both panels, regions of the tissue containing nodose and jugular neurons have been highlighted, matching the output from RCTD. Beads were assigned nodose (purple colours) and jugular (blue) neuronal subtypes by RCTD. **(E)** Neighbourhood analysis on the RCTD output, including the cell types assigned to singlet and multiplet beads. We identified 5 distinct neighbourhood types. Neighbourhoods were labelled based on the cell types which co-occur in the regions. Neighbourhoods highlighted in blue predominantly contain glial cell types, NGN/Glia-like neighbourhoods are highlighted in pink, orange and green, and the JGN/Glia-like neighbourhood is indicated in purple (see Figure S4 legend for more details on expression patterns in each neighbourhood). **(F-G)** *Phox2b* is a marker highly expressed in nodose neurons with only low expression in jugular neurons. The cells expressing *Phox2b* were coloured based on the same scale bar. **(F)** log-normalised expression of *Phox2b* in 1 tile of the spatial transcriptomics dataset. **(G)** log-normalised expression of *Phox2b* in the single cell atlas. *Phox2b* expression is enriched in nodose ganglion neuronal clusters. **(H)** Diagram highlighting the orientation of the nodose ganglion sections in the mouse. **(I)** Heatmap showing scaled expression of marker genes in nodose ganglion neuronal clusters. **(J)** UMAP plots of nodose and jugular neurons, coloured by neurotransmitter (left) and neuropeptide (right) assignment.

In mice, the nodose and jugular ganglia merge to form the vagal ganglia^21^. Our analysis identified a total of 53 clusters, and we annotated them based on cell type and three marker genes with the highest cluster specificity (Figure 1A, I, Table S1, S2). We identified 22 nodose and 5 jugular neuronal clusters, 3 glial, 8 satellite glial and 6 myelinating glial cell clusters, 3 endothelial, 4 fibroblast and 2 hemopoietic cell clusters (Figure 1A, Figure S2, Table S3). To validate our clustering strategy with other datasets included in NodoMap, we compared our clustering with that reported by Kupari et al^14^ and Buchanan et al^28^, and found that while we had slightly higher nodose neuronal granularity, cell type grouping was well preserved across the studies (Figure S3). We also identified more non-neuronal clusters than Kupari, highlighting improved cell type segregation through integration of multiple datasets and higher cell count. Interestingly, the number of jugular neuronal clusters remained similar, suggesting limited heterogeneity. In addition to snRNASeq we performed single-cell resolution (10µm) spatial transcriptomics (Curio Seeker platform^36^) on a total of 9 nodose ganglia sections from the left and right ganglia. Integration of snRNASeq and spatial transcriptomics was performed using the Robust Cell Type Decomposition (RCTD)^37^ function in Seurat, which allowed us to map the majority of the snRNASeq clusters onto both the left and right nodose ganglia (Figure 1B-D)^37^. As expected, expression of the nodose ganglia neuronal marker *Phox2b* was able to distinguish between nodose and jugular neurons in the snRNASeq dataset^38^, and was also found in regions where nodose neuronal clusters were mapped in the spatial transcriptomics dataset (Figure 1F-G). We additionally saw some low-level *Phox2b* expression in myelinating glial cells, satellite glial cells and other glial cell populations^38^ in our spatial transcriptomics dataset. NGN5, which is characterised by *Rtn1, Ngfr* and *Clu*, was the neuronal nodose cluster mapped to the spatial dataset most frequently.

The spatial relationships between different nodose ganglia cell types have not previously been comprehensively evaluated. We carried out cellular neighbourhood analyses (See Methods)^39^ to investigate whether specific multicellular combinations were more likely to be found together, indicating structural organisation within the ganglia. We identified five types of neighbourhoods from the spatial transcriptomics data: one comprised predominantly of non-neuronal cells (EC3, FB4, EC2, HC2 and NGN5), one comprised of jugular ganglia neurons and glial-like cells (JGN1 JGN3 JGN4 and non-neuronal cells like GB3 MGC4 and MGC2, among others), and three comprised of different nodose ganglia neuronal populations and glial-like cells (Figure 1E and Figure S4). These neighbourhoods were equally present in both left and right nodose ganglia (data not presented).

We also characterised neuronal subtypes through their expression of neurotransmitter and neuropeptide associated genes. As expected, the vast majority of nodose neurons were glutamatergic, with specific clusters expressing distinct elements of the glutamatergic molecular machinery. Of note, while most nodose neuronal types expressed only the vGLUT2 (*Slc17a6*) vesicular glutamate transporter, two nodose clusters and one jugular neuronal cluster expressed both vGLUT1 (*Slc17a7*) and vGLUT2 (Figure 1J; Figure S4B), suggesting specialised differences in the synapses associated with these neurons, and differences in the probability of glutamatergic release from these neurons in comparison to other neuronal populations^40^. A number of other glutamatergic clusters were also found to be dopaminergic (Figure 1J), and NGN20, marked by the expression of *Slc6a2, Npy and Hand2*, lacked expression of glutamatergic genes but expressed *Dbh, Ddc* and *Th* transcripts. *Dbh* encodes dopamine beta-carboxylase, an enzyme which converts dopamine into noradrenaline, suggesting that this neuronal population is likely noradrenergic (and contains the necessary molecular machinery for dopamine synthesis). This cluster was previously identified as representing ‘sympathetic neurons’ by Kupari et al^14^ (Figure S3). Nodose neurons also expressed genes encoding different combinations of neuropeptides (Figure 1J, Figure S4B), including *Tac1, Npy, Nts, Cartpt and Adcyap1* (Figure 1J, Figure S34B). *Npy* expression was enriched in NGN20.Slc6a2/Npy/Hand2, and this cluster also expressed *Gal*. *Vip* transcripts were found in NGN21, NGN4 and NGN14. (Figure 1J). Regarding notable membrane sensors and channels, a cluster of neurons expressed the Cck A receptor (*Cckar*) (NGN5), and other discrete clusters expressed *Trpa1* (NGN9), *Slc18a3* (NGN13) and *Olfr78* (NGN22).

In addition to classifying neurons by expression of neurotransmitters and neuropeptides, using previously reported information^14,15,30^, we classified each of our neuronal clusters based on the type of expressed sodium channels, myelination level, the sensor type, and the organ each cluster innervated (Figure S5). Of the 27 neuronal clusters, 10 were identified as expressing predominantly Nav1.1 channels, and 15 as expressing predominantly Nav1.8 channels, with 2 clusters expressing similar levels of both channels (Figure S5A). Myelination affects the speed at which action potentials travel along axons, and thus affects the speed of signalling to the brain. Only 5 neuronal populations were found to be myelinated (JGN3, NGN15, NGN13, NGN11, NGN22), as determined by expression of *Nefh*, *Cntn1*, *Cntnap1*, and *Ncam1*. Twelve neuronal clusters were unmyelinated and 10 were lightly myelinated (Figure S5B). Neurons in the nodose and jugular ganglia sense mechanical and nociception information from the periphery and relay this information to the CNS. Altogether, 10/27 neuron clusters were identified as mechanosensors, 11/27 as nocisensors, and 6 were identified as a mix of the two (Figure S5C).

Finally, nodose and jugular neurons communicate information from different organs to the brain. Using transcriptomic signatures, we predicted the organ where each neuronal population innervated. Of the 5 jugular neuronal clusters, JGN3 transduces information from the gut, JGN5 senses information from the lungs, and the other 3 jugular clusters displayed a broad projection pattern. With regards to nodose neurons, NGN8 innervated the pancreas, NGN16 the heart, NGN22 the jejunum/ileum, NGN13 the duodenum and NGN5 the stomach (Figure S5D). Previously reported gene markers for vagal afferents innervating the gut ^15^ were also expressed in another 8 nodose neuronal clusters, which we also classed as projecting to the gut. An additional 9 nodose neuronal clusters expressed gut marker genes and also organ projection marker genes to one or more other organs, and were thus classed as having broad projections. To assess whether different classifications of neurons were organised spatially into distinct neighbourhoods, we transferred the sodium channel/fibre type/sensor type/organ projection labels to the spatial transcriptomics dataset and re-ran neighbourhood analysis (Figures S6-9). Interestingly, despite suggestions that the left and right ganglia innervate different organs to higher or lesser degrees^18,28,29^, we observed no differences between ganglia in the pattern of neurons as classified by organ innervation.

Afferent vagal neurons synapse with neurons of the dorsal vagal complex (DVC) in the hindbrain, predominantly in the nucleus tractus solitarius (NTS) and the area postrema (AP)^18,41^. To identify possible inter-neuronal communications between the nodose/jugular ganglia neurons and neuronal populations within the hindbrain, we employed CellChat^42^, to explore possible ligand-receptor interactions between NodoMap and a previously published single-cell transcriptomic dataset of the mouse hindbrain^43^. We identified a total of 57 potentially enriched pathways signalling in the biologically relevant direction (nodose/jugular to hindbrain neurons; Table S13). To categorise signalling pathways based on how many neuron types they are signalling from/to, we classified pathways as having a ‘many’ relationship if the number of source or target clusters was greater than 25% of the total clusters (and if less than 25% then this was a ‘few’ relationship). In this instance, we identified 28 pathways with a many-to-many nodose to hindbrain neuronal connectivity, 15 pathways with a few-to-many relationship, 6 with many-to-few and 8 pathways with a few-to-few nodose to hindbrain neuronal connectivity.

As expected, a common signalling pathway from all nodose neuronal clusters to all hindbrain neuronal clusters was glutamatergic signalling, with no GABAergic signalling identified between the two regions^44–46^. To focus on nodose to AP/NTS specific signalling pathways, we identified six neuronal clusters in the hindbrain dataset that were highly likely to originate from the AP or NTS, based on expression of AP/NTS specific marker genes (see methods). One pathway of particular interest was TAC signalling, which highlighted 9 source clusters from the nodose/jugular neurons, and 4 target clusters in the hindbrain neuronal dataset (many-to-few, Figure S10A). Four of the source clusters were nodose neuronal clusters NGN11, NGN13, NGN15 and NGN17, all of which potentially signal to a specific neuronal population in the DVC: HB_NE_Gcg/Prlr, likely to be GLP-1 producing neurons known to be involved in modulation of appetite^47^. All four of these nodose clusters were predicted to project from either the duodenum specifically or the gut more generally (Figure S5D), highlighting a possible gastrointestinal-nodose-DVC signalling pathway via *Tac1/Tac1r*^48,49^. Additionally, energy homeostasis associated gene (*Enho*) signalling was identified with a few-to-many signalling relationship (Figure S10B). *Enho* encodes the small peptide Adropin, reported to bind and signal via the G-protein coupled receptor 19 (*Gpr19*)^50,51^. All nodose neuronal clusters expressed at least some levels of *Enho,* with potential target clusters in the hindbrain including HB_NE_Tbx20/Prph, thought to be NTS specific due to its expression of transcription factor *Phox2b*. A list of all identified signalling pathways, their ligands and receptors, source and target clusters, and their classifications is shown in Table S13.

The vagus nerve plays a crucial role in the gut-brain axis, regulating feeding behaviour in response to food ingestion^13,15,52^. Fasting has previously been reported to alter the expression of specific genes associated with energy homeostasis in the nodose ganglia^22,25,53,54^, but the changes occurring in individual neuronal types are largely unknown. We performed snRNASeq on the nodose ganglia of mice fasted overnight and compared the expression profile to that from *ad libitum* fed mice, using data from 680 nuclei from overnight fasted mice and 1275 from ad libitum fed animals. These nuclei were distributed across nearly all of the clusters we identified across the five datasets (51 out of 53 clusters, with no nuclei found from the MGC6 and NGN21 clusters) (Figure 2A-B). As expected, fasting did not alter cell cluster identity, but differential gene expression analysis within each nodose neuronal cluster identified a number of genes upregulated or downregulated in specific nodose neuronal populations by fasting (Figure 2C; Table S4). NGN2, NGN6 NGN9 and NGN11 appeared particularly transcriptionally sensitive to fasting. NGN2 and NGN11 displayed the greatest number of genes downregulated and NGN9 and NGN6 had the greatest number upregulated in response to fasting. These changes included the upregulation by fasting of the gene encoding Amyloid-like protein 1 in the NGN2 cluster, which has been implicated in the modulation of glucose and insulin homeostasis^55^, and the downregulation of the gene encoding the Metabotropic glutamate receptor 8 in NGN11 (Figure 2D-G). Ingenuity Pathway Analysis of the altered expression profiles in these four neuronal clusters found diverse signalling and metabolic pathways were altered in specific clusters, including the downregulation of nutrient and energy sensitive pathways such as mTOR signalling and the response to amino acid deficiency in NGN6, and the up or downregulation of AMPK signalling in different clusters (Figure 2H, Table S6).

**Figure 2:**
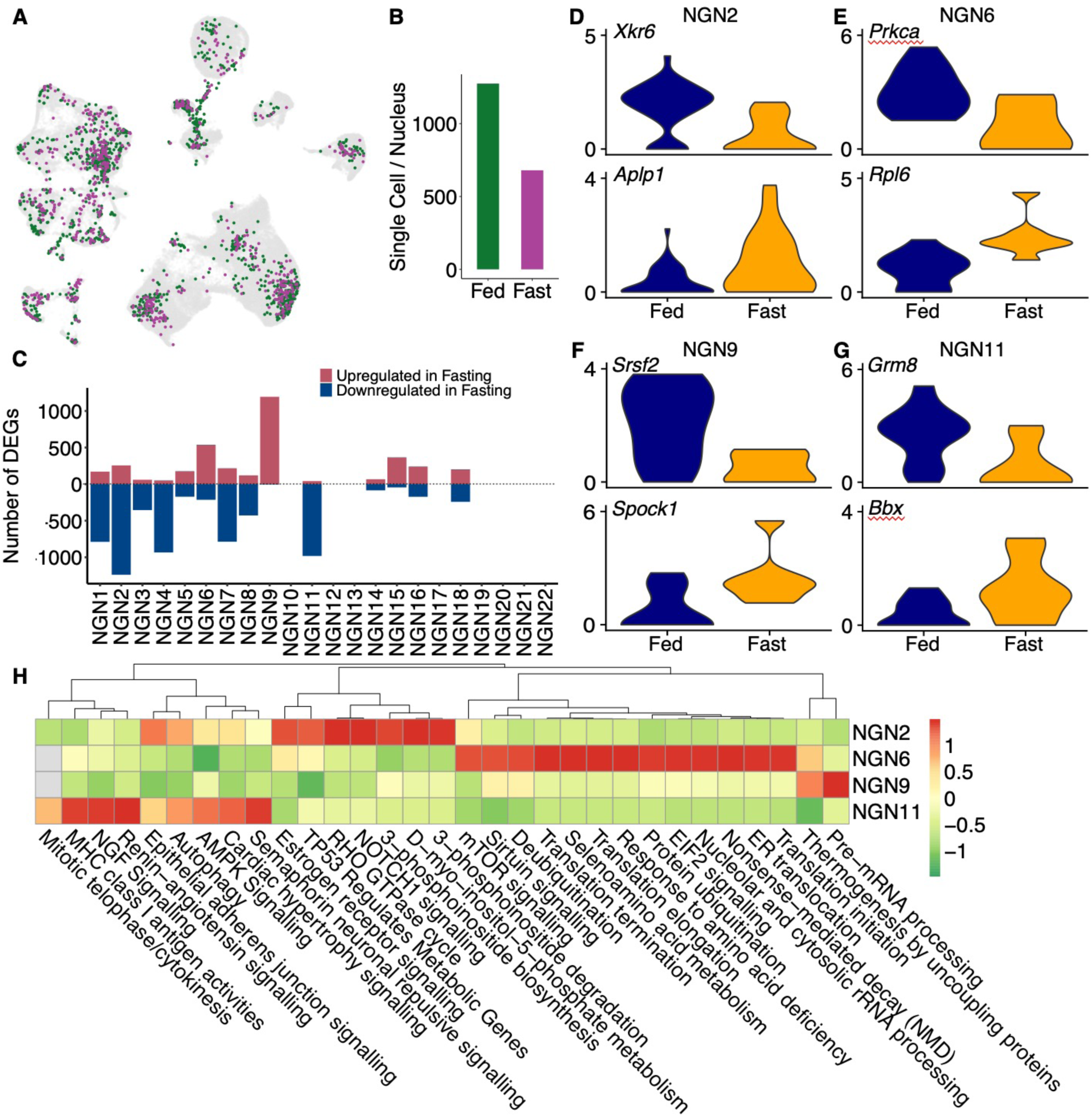
The effects of fasting on nodose ganglion neuronal gene expression. **(A)** A UMAP plot with cells from *ad libitum* fed (green) and overnight fasted mice (pink) highlighted. **(B)** The number of nuclei originating from *ad libitum* fed and overnight fasted mice. **(C)** The number of significantly upregulated (red) and downregulated (blue) genes in each nodose neuronal cluster in response to overnight fasting (P < 0.05). **(D-G)** The top upregulated and downregulated gene in NGN2, NGN6 NGN9 and NGN11 in fasted animals, selected based on the specificity score and their expression levels (see methods) **(H)** The top 10 pathways that were enriched in the top 4 transcriptionally sensitive nodose neuronal clusters (see methods).

Current literature suggests functional differences in the left and right nodose ganglia, with, for example, the left nodose ganglion thought to be more involved in distension-induced satiety, and the right nodose ganglion having a greater role in food preference^18^. Differential gene expression was analysed using in-house data and Buchanan et al^28^, totalling 11,282 cells or nuclei from the left nodose and 8565 from the right nodose ganglia (Figure 3 A, B), distributed across all the 53 clusters (Figure 3A). Similar numbers of cell clusters and distributions of the five neighbourhoods were observed in both left and right ganglia (Figure 3D, F, H, J). However, all clusters showed differential expression of specific genes between left and right ganglia (Figure 3K; Table S5). In particular, clusters NGN5, NGN10, NGN17, NGN19 had 942, 1900, 1671 and 1184 significantly differentially expressed genes (DEGs), respectively (Figure 3K-O). Pathway analysis on the significantly DEGs in these nodose neuronal clusters identified a number of signalling pathways enriched in left or right nodose neurons. For example, glutamatergic receptor signalling pathways were enriched in right nodose neurons in the NGN5 and NGN10 clusters, and estrogen receptor and opioid signalling were enriched in right nodose neurons in NGN5. Interestingly, given that the right ganglia may be considered more nutrient sensitive, mTOR signalling was also enriched in NGN5 neurons from the right ganglia (Figure 3P, Table S7).

**Figure 3:**
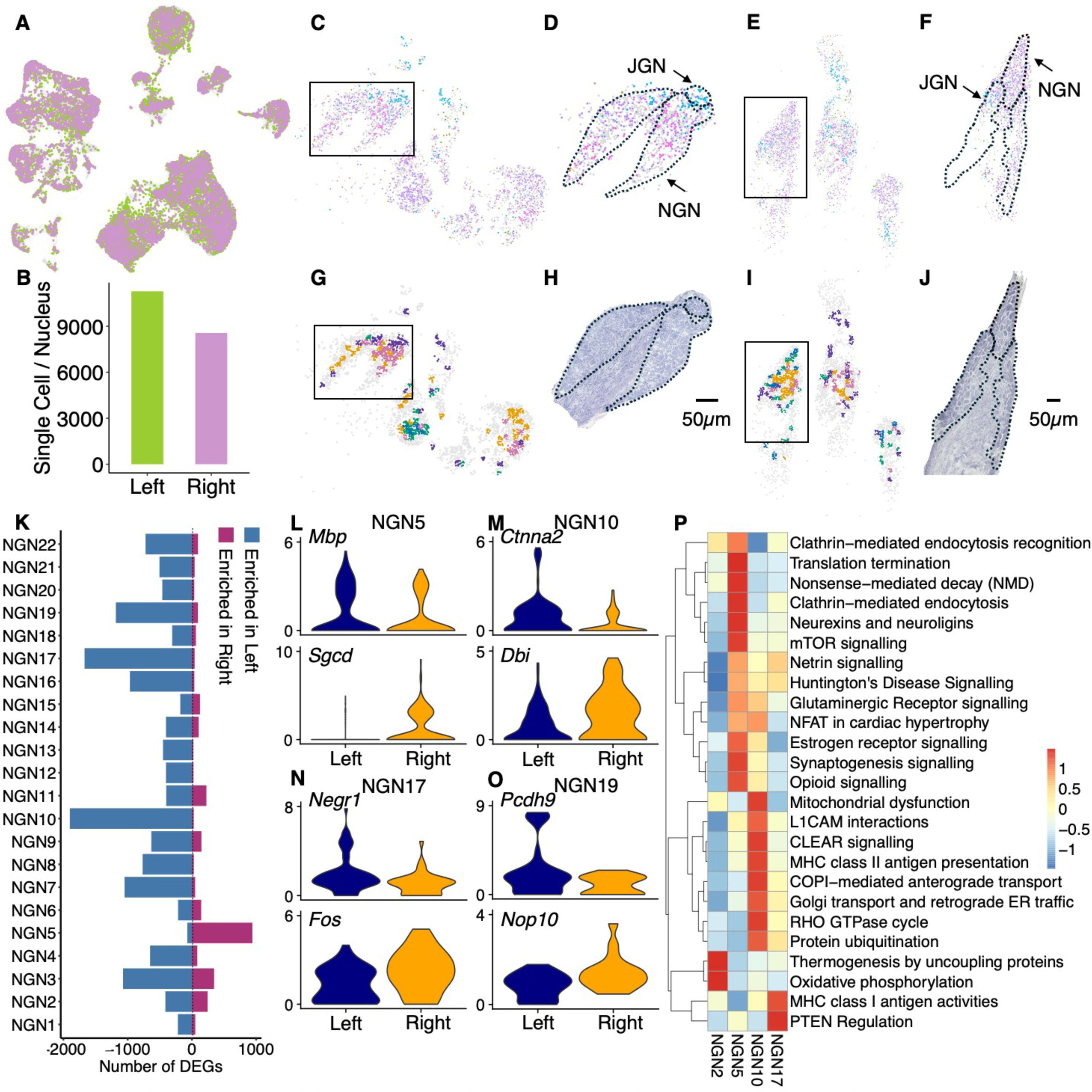
Differences in gene expression between the left and right nodose ganglia. **(A)** A UMAP plot highlighting cells which originated from left nodose ganglia (green) and right nodose ganglia (lilac). **(B)** The number of cells from left or right nodose ganglia within the dataset. **(C)** RCTD mapping of snRNASeq clusters on 3 left nodose ganglion sections on 1 tile, with panel **(D)** showing a zoomed-in image of the highlighted region in C. Regions containing nodose or jugular neurons are highlighted. **(E)** RCTD mapping of snRNASeq clusters on 3 right nodose ganglion sections on 1 tile, with panel **(F)** showing a zoomed-in image of the highlighted region in E. Regions containing nodose or jugular neurons are highlighted. **(G)** Neighbourhoods identified in the left nodose ganglia, with NGN/Glia like neighbourhoods highlighted in pink, orange and green, JGN/Glia-like neighbourhoods highlighted in purple and Glia-like neighbourhoods highlighted in blue. **(H)** Haematoxylin stained near adjacent tissue section of the nodose ganglia highlighted in the box in C and D. NGN and JGN regions highlighted by dotted lines. **(I)** Neighbourhoods identified in the right nodose ganglia, with NGN/Glia like neighbourhoods highlighted in pink, orange and green, JGN/Glia-like neighbourhoods highlighted in purple and Glia-like neighbourhoods highlighted in blue. **(J)** Haematoxylin stained near adjacent tissue section of the nodose ganglia highlighted in the box in E and F. NGN and JGN regions highlighted by dotted lines. **(K)** DEG analysis comparing left vs right in nodose neuronal clusters. Some clusters displayed particularly high numbers of DEGs (P < 0.05). NGN5, NGN10, NGN17, NGN19 had 942, 1900, 1671 and 1184 significantly differentially expressed genes, respectively, with **(L-O)** showing the top genes enriched in left/right in each of these 4 clusters. **(P)** Pathway analysis on the DEGs (P < 0.05) in the top 4 transcriptionally sensitive nodose neuronal clusters.

In accord with previous analyses, jugular neurons organised into fewer clusters than nodose neurons^14^. Jugular neurons were identified by enriched expression of Prdm12^14^ and formed five clusters distinct from the nodose clusters (Figure 4A, B) and defined by enrichment of individual genes (Figure 4C). Interestingly, JGN5 expressed the cold and menthol receptor Trpm8 thought to be involved in oesophageal vagal sensory signalling^56^. Jugular neuron clusters also showed a number of DEGs in response to fasting, and between the left or right ganglia (Figure 4D-I), though the numbers of DEGs were lower than in nodose neurons. The JGN1 and JGN5 clusters showed the highest number of significantly downregulated (890) and upregulated (612) DEGs in response to fasting, respectively. The nutrient sensitive and metabolism pathways altered in response to fasting in JGN clusters were similar to those observed in NGN clusters (Table S15). For example, the response to amino acid deficiency pathway was enriched in JGN1 and the regulation of lipid metabolism by PPARa pathway was upregulated in JGN5. JGN1 also had the highest number (451) of DEGs enriched in the left ganglia and JGN3 had the highest number (185) in the right ganglia. Oxytocin and glutaminergic receptor signalling pathways were downregulated in left jugular neurons in JGN1, and nutrient sensitive pathways such as the response to amino acid deficiency enriched in right jugular neurons in JGN3 (Table S16).

**Figure 4:**
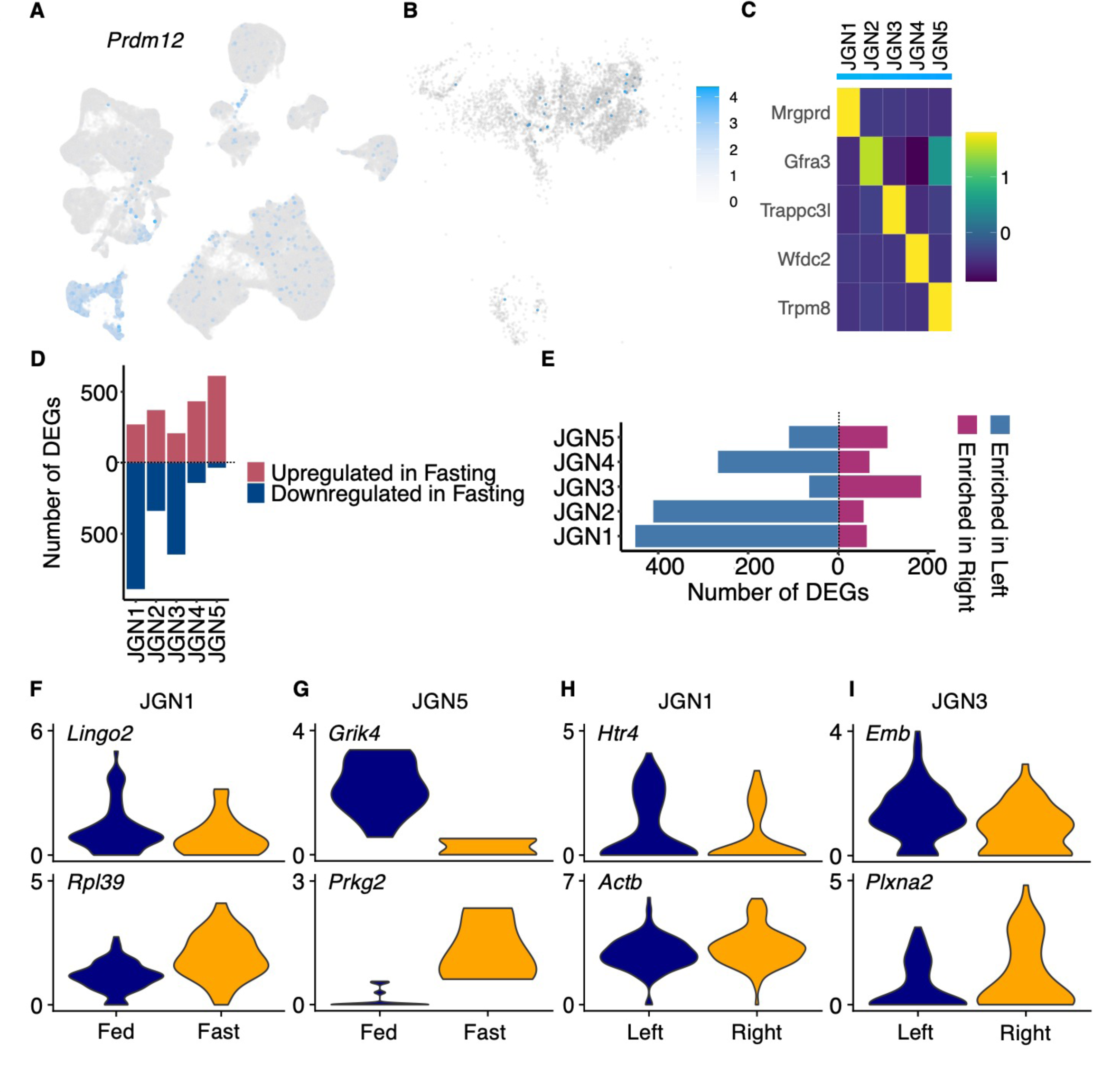
Overview of Jugular neurons. **(A)** Log-normalised expression of *Prdm12* in NodoMap. **(B)** Log-normalised expression of *Prdm12* in spatial transcriptomics dataset. Cells were coloured by the same expression level scale bar as A. **(C)** Heatmap showing scaled expression of five marker genes in jugular ganglion neuronal clusters **(D-E)** Graph showing the number of statistically significantly DEGs (P < 0.05) in each jugular neuronal cluster **(D)** when comparing overnight fasting to *ad libitum* fed (red: upregulated in fasting, blue: downregulated in fasting), **(E)** and left vs right (red: enriched in right nodose ganglia, blue: enriched in left nodose ganglia). **(F-I)** Violin plots showing log normalised expression of top DEGs in (F-G) fed and fasted animals, and in (H-I) left vs right nodose ganglia.

Twenty six clusters of non-neuronal cell types were identified in both the snRNASeq and spatial transcriptomics datasets by the presence of the marker genes *Emcn* (endothelial cells), *Ebf2* (Fibroblasts), *Ptprc* (Hematopoietic cells), and *Mpz* (Myelinated glial cells) and *Fabp7* (Satellite glial cells)^14,57–59^ (Figure 5A-K). Again, specific cell types showed differential gene expression between fed and fasted states, and between left and right ganglia (Figure 5L-Q), though the numbers of genes with altered expression was typically lower than the numbers altered in the neuronal clusters. Cluster MGC1 had the highest number of fasting downregulated DEGs (258) and SGC1 had the highest number of fasting upregulated DEGs (357). Intracellular transduction and myelination signalling pathways were either downregulated or upregulated by fasting across different non-neuronal clusters (Table S15). Comparing left and right ganglia, FB3 had the highest number (1024) of enriched DEGs in the left ganglia, and FB4 the highest number (815) enriched in the right ganglia. Interestingly, estrogen receptor and insulin-like growth factor 1 signalling were both downregulated in the left ganglia of FB3. In contrast, both estrogen receptor and cholecystokinin/gastrin-mediated signalling were enriched in the right ganglia of FB4 (Table S16).

**Figure 5:**
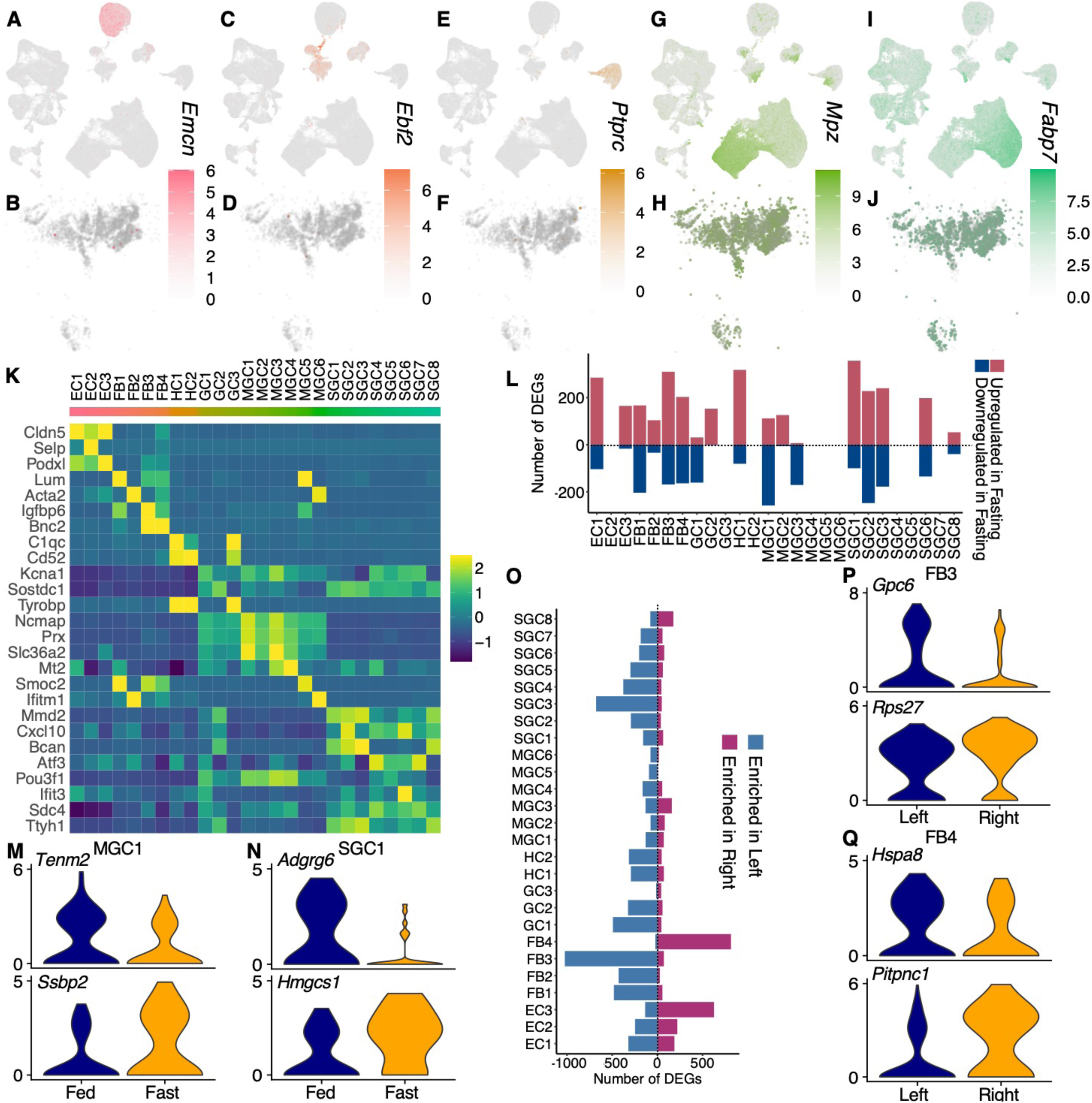
Overview of non-neuronal cells. **(A, C, E, G, I)** Log-normalised expression of non-neuronal cell markers in single cell atlas and **(B, D, F, H, J)** a tile of the spatial transcriptomics dataset of 3 left nodose ganglia. For each marker gene, the cells were coloured by the same expression scale bar. (A, B: *Emcn*: Endothelial cells (EC); C, D: *Ebf2*: Fibroblasts (FB); E, F *Ptprc*: Hematopoietic cells (HC); G, H: *Mpz*; Myelinated glial cells (MGC); I, J: *Fabp7*: Satellite glial cells (SGC). If both MGC and SGC cell markers were presented in the clusters, they were labelled as glial cells (GC)). **(K)** Heatmap with scaled expression of marker genes in non-neuronal clusters. **(L)** Number of significantly DEGs in fasting across all non-neuronal clusters (P < 0.05). **(M-N)** Violin plots showing top DEGs in the fasted state **(O)** Number of significantly DEGs comparing left and right nodose ganglia across the non-neuronal clusters (P < 0.05). **(P-Q)** Violin plots with marker genes enriched in left or right nodose ganglia from FB3 (P) or FB4 (Q) cluster.

## Discussion

We have harmonised multiple single cell/nucleus RNASeq datasets of the mouse nodose ganglia and aligned them with spatial transcriptomics data to create a tool for researchers interested in the physiological and pathophysiological roles of the vagus nerve. Research into vagal signalling has advanced significantly in the last decade^15,17,45,46,52,60^, helped greatly by new molecular approaches, and we hope that NodoMap will facilitate further understanding of this major gut-brain pathway. Harmonising data from multiple sources is likely to improve the resolution of cell classification, particularly for rarer populations. Figure S3 shows we have refined resolution when comparing nodose ganglion neuronal clusters (e.g. NG15 and NG16)^14^.

The approach we have taken here is similar to that we have previously employed for the mouse^34^ and human hypothalamus^61^. However, even though we have not had opportunity to perform similar analyses on human tissue, evidence suggests that human nodose ganglia have similar characteristics to those in the mouse, including expression of key genes and overlapping functional pathways^62^.

CellChat analysis identified a number of potential signalling pathways between the vagus and the hindbrain. Although a number of specific vagal-brainstem pathways have been characterised^45,63^, there are likely to be many more, with the role of, for example, 2-AG and prostaglandin^64^ synthesised in nodose neurons having been little examined, and the roles of neuropeptide signalling from the vagus to the hindbrain also remaining largely unknown. While the CellChat analysis likely overestimates possible connections, as the hindbrain dataset used will include many cells not innervated by vagal neurons, this analysis suggests new pathways to investigate using functional studies. Future experiments using spatial transcriptomics analysis of the DVC linked to tracing of vagal terminals may further help to focus further projects on the most promising circuits.

In summary, NodoMap provides a rich resource for researchers studying vagal signalling. It is intended as a dynamic tool, regularly updated as new data comes to light, and to be compared with any future equivalent analysis of the human nodose ganglia. In combination with the necessary functional studies, NodoMap will thus hopefully support the discovery of new approaches to exploiting vagal signalling in a clinical context.

## Acknowledgments

The Section of Endocrinology and Investigative Medicine is funded by grants from the MRC, BBSRC, NIHR and is supported by the NIHR Biomedical Research Centre Funding Scheme and the NIHR/Imperial Clinical Research Facility. The views expressed are those of the authors and not necessarily those of the MRC, BBSRC, the NHS, the NIHR or the Department of Health. KGM is supported by Diabetes UK (18/0005886, 20/0006295), the BBSRC (BB/W001497/1, BB/X017273/1), the MRC (MR/Y013980/1) and the Wellcome Trust (310835/Z/24/Z). SC was supported by the British Society of Neuroendocrinology. BYHL and GSHY are supported by BBSRC Project Grant (BB/S017593/1) and the MRC Metabolic Diseases Unit (MC_UU_00014/1). Next-generation sequencing was performed at the IMS Genomics and Bioinformatics Core supported by the MRC (MC_UU_00014/5, RGAG/542 MC_UU_00039) and the Wellcome (208363/Z/17/Z, RGAG/546 226800/Z/22/Z), and the Cancer Research UK Cambridge Institute Genomics Core.

## Declaration of interests

BYHL provides remunerated consultancy for Nuntius Therapeutics. GSHY receives grant funding from Novo Nordisk and Amgen Inc; he also consults for both Novo Nordisk and Eli Lilly and Company. The other authors declare no competing interests.

## STAR METHODS

### EXPERIMENTAL MODEL AND STUDY PARTICIPANT DETAILS

A total of twenty-six 7-week-old male C57BL/6J mice (Charles River, UK; RRID: IMSR_JAX:000664; twenty for snRNASeq and 6 for ST) were maintained on a 12hr light/dark cycle under a constant temperature (21-23°C) with free access to standard chow and water. Mice were group housed in ventilated cages and pathogen free facilities. All animals were acclimatised to the experimental procedures and randomised by body weight. For snRNASeq experiments, ten mice were fasted overnight for 16-hours, and another ten mice were *ad libitum* fed prior to nodose ganglia extraction. Mice were randomly assigned to experimental groups by bodyweight.

All animal studies were performed according to the UK Home Office Animals Scientific Procedures Act 1986 (Project License No. PD75F462C) and approved by the Central Biomedical Services unit at the Hammersmith Campus, Imperial College London.

## METHOD DETAILS

### Nuclei isolation

Mice were euthanized by decapitation at the same time of day and the nodose ganglia were immediately extracted and snap frozen on dry ice pooling overnight-fasted left, overnight-fasted right, *ad libitum* fed left, and *ad libitum* fed right nodose ganglia into four separate 1.5mL Eppendorf tubes. The pooled nodose ganglia samples were stored at -80°C before until nuclei isolation.

Each sample was separately homogenized in 500µL homogenization buffer (100µM DTT, 0.1% Triton X-100, 2x protease inhibitor, 0.4U/µL RNasin, 0.2U/µL Superase.In, 1µl/mL Draq5, and nuclei isolation medium (250mM sucrose, 25mM KCl, 5mM MgCl2, 10mM Tris buffer (pH 8.0) in nuclease-free water) in a 2mL dounce homogenizer on ice using 10 strokes with pestle A and 10 strokes with pestle B. Samples were transferred to separate 1.5mL Eppendorf tubes and centrifuged at 900 r.c.f at 4°C for 10 minutes. The supernatant was carefully removed and pellets were each thoroughly resuspended in a 1:1 mixture of homogenization buffer and 50% Optiprep diluted in iodixanol dilution media (total volume 450µL per sample, final Optiprep concentration was 25%; 250mM sucrose, 150mM KCl, 30mM MgCl2, 60mM Tris buffer (pH 8.0), and nuclease-free water). These suspensions were then each carefully layered on top of separate 450µL 29% Optiprep diluted in iodixanol dilution medium in 2mL Eppendorf tubes and were centrifuged at 13,500 r.c.f, at 4°C for 20 minutes to separate the nuclei and debris. The supernatant was removed, leaving the nuclear pellet. Pellets were resuspended in 1mL PBS containing 0.04% BSA.

### Molecular tagging of nuclei

Resuspended samples were centrifuged at 900 r.c.f for 10 minutes at 4°C and supernatant was removed. Samples were resuspended in 100µL of different cell multiplexing oligo (CMO; from the 10X Genomics 3’ CellPlex kit) and incubated for 5 minutes at room temperature. The CMO is a feature barcode oligonucleotide connected to a lipid. After incubation, 1.9mL resuspension buffer (PBS with 1% BSA and 0.4U/µL RNasin) was added to each sample, and the mixture centrifuged at 900 r.c.f for 10 minutes, and then the supernatant was removed. Another 1mL resuspension buffer was added, the mixture centrifuged at 900 r.c.f for 4 minutes and then the supernatant removed, to remove any unbound CMO. Nuclei pellets were resuspended in 100µL resuspension buffer each and then combined. The pooled sample was centrifuged at 900 r.c.f for 10 minutes, the top 50µL of supernatant removed, and the pellet resuspended in the rest of the supernatant.

### snRNASeq library preparation

The snRNASeq library was prepared according to the Chromium Next GEM Single Cell 3□ Reagent Kits v3.1 (Dual Index) with Feature Barcode technology for Cell Multiplexing protocol. Nuclei were loaded onto the Chromium Chip G along with master mix, gel beads and partitioning oil and the chip was placed into the Chromium controller. In the machine, individual nuclei are encapsulated inside partitioning oil along with a single Gel Bead to form GEMs. Each Gel Bead contains oligonucleotides which consist of cell barcodes and unique molecular identifiers (UMI) and a poly(dT) tail which capture mRNA transcripts. A cDNA library is generated through reverse transcription; bead clean up and pre-amplification PCR. The cDNA library is then fragmented enzymatically amplified once more to create the final RNA sequencing library. The snRNASeq library was sequenced on a NovaSeq 6000 (Illumina, USA) at a minimum sequencing depth of 30,000 reads per nucleus.

### Tissue preparation for Spatial Transcriptomics

After cervical dislocation, the left and right nodose ganglia from six mice were dissected and snap frozen on dry ice. Three nodose ganglia in the same orientation were placed on a single disposable base mould (Leica Biosystems, UK) and embedded by adding chilled bubble free optimal cutting temperature compound (Sakura 4583 Tissue-Tek O.C.T. Compound, USA). The samples, comprising 6 left nodose ganglia and 3 right nodose ganglia in total, were stored at -80°C until sectioned on a cryostat at -20°C into 10µm sections (Leica CM1950, Leica Biosystems, UK). The 10µm section was mounted to a pre chilled 3mm by 3mm Curio Seeker tile slide (Curio Bioscience, USA) for high-resolution spatial transcriptomics.

The 10µm sections sliced before and after the section that was mounted on the Curio Seeker tile were mounted on Superfrost Plus Adhesion Microscope Slides (Epredia, UK) for haematoxylin (Sigma-Aldrich, MHS32) staining to visualise the sample morphology.

### Spatial Transcriptomics library preparation

The cDNA library was generated according to the Curio Seeker Spatial Mapping Kit (3×3) - User Manual for Fresh Frozen Tissues. Briefly, the tile with tissue mounted on was placed in an Eppendorf containing hybridization mix and incubated to allow for mRNA to hybridize to the barcoded poly (dT) oligos on the lawn of 10µm beads on the tile. Next, reverse transcription occurred to generate double stranded cDNA and the tissues were digested to release the beads into solution. cDNA was amplified, purified and quantified. Next, the Illumina sequencing platform compatible libraries were prepared by using the Nextera XT DNA Library Preparation Kit (Illumina, USA). The library was sequenced on Novaseq 6000 (Illumina, USA, Read1 = 50bp, Read2 =50bp, Index1 and Index2 = 8bp).

## QUANTIFICATION AND STATISTICAL ANALYSIS

### Dataset downloads

The droplet-based 10X single cell sequence reads of mouse nodose ganglia from 4 publicly available datasets^14,15,28,30^ were downloaded using FASTQ-dump from the sequence read archive prefetch toolkit (NCBI SRA version 3.0.0) (GSE124312, GSE138651, GSE185173, GSE192987).

### Sequence alignment

Data was aligned to the mouse transcriptome (GRCm38 mm10, Ensembl 100) using the cellranger count function (version 6.0.1, 10X Genomics, USA). The same reference was used for all datasets (both accessed online, and data created in-house). Sample demultiplexing was performed as follows: Mapping of trimmed 15bp (fastx_trimmer 0.013) 10X Nextera CMO reads (R2) to the reference CMO sequences provided by 10X Genomics using bwa’s mem command, version 0.7.17), The aligned bam file was then annotated with cell barcodes (XC) and unique molecular identifiers (XM) extracted from R1 (fastxtrimmer 0.0.13) usingfgbio AnnotateBamWithUmis command (version 1.5.1). Reads were then filtered to only those annotated with valid cell barcodes. Only cells with more than 150 valid CMO reads were retained for downstream analyses. The sample assignment and CMO counts are listed in Supplementary Table S17 and S18.

### Dataset quality control

Scater (Version 1.32.0)^65^, DropletUtils (Version 1.20.0)^66,67^, scDblFinder (Version 1.11.4)^68^, and Seurat (Version 4.3.1)^69^ packages were used for data pre-processing: removing low quality cells and identifying and removing doublets, prior to integration. Datasets were filtered on an individual basis to identify and remove low quality cells based on a cut off for the number of genes, UMIs and percentage of mitochondrial genes per dataset. The cut-offs used for each dataset can be found in Table S10. Doublets were identified and removed using scDblFinder. Metadata was added to each dataset, including the source of datasets, the age and the strain of mice, the position of nodose ganglia (left or right), and the nutritional status of the mice.

### Batch correction and clustering

Datasets were integrated according to the seurat integration pipeline for SCTranform normalised data. The pre-processed datasets were imported as a list, normalised using SCTransform (Version 0.3.4)^70^. The highly variable gene list was selected by ranking each gene based on the number of datasets it was deemed to be highly variable in. The integration method was iterative pairwise integration. After integrating all nodose ganglia cells/nuclei, batch-corrected datasets were clustered using the Louvain algorithm (FindNeighbors and FindClusters from Seurat Version 4.3.1^69^) and a quality control performed to check the count and feature reads of each cluster. Any clusters with obviously low counts/genes per cell were removed as low-quality clusters. The final integrated nodose single cell/nucleus data had a total of 55,399 genes and 108,482 cells or nuclei.

To optimize the clustering of the dataset, we clustered by computing the k-nearest neighbours based on a set of different number of principal components (PC = 30 to 60, incremental increase by 10), and ran Louvain clustering at differing resolutions (0.6 to 2, incremental increase by 0.2). The optimal number of clusters was selected based on the average silhouette width^71^ (ASW) of overall and *Phox2b* expressed clusters. *Phox2b* is a marker gene of nodose ganglia neurons (NGN)^14^. ASW measures the distance between cells in one cluster to the cells in the neighbouring clusters with a range of [-1, 1]. The closer of ASW to +1, the more distant the cells are from the neighbouring clusters, and the negative ASW suggests the cells are assigned to the wrong clusters. In summary, 1) 25,000 cells were randomly selected from the dataset analysed for each PC and resolution; 2) their coordinates from 1 to 50 PCs were used to calculate the distances were calculated using the first 50 PCs between cells to generate a dissimilarity matrix. Silhouette widths were calculated based on the dissimilarity of cells using silhouette function from Cluster (Version 2.1.6)^72^; 3) step 1) and 2) were repeated five times and all silhouette widths averaged to get the overall ASW; 4) for cells calculated from the same PC and resolution, their overall ASWs were aggregated and averaged. The averaged overall ASW of each PC and resolution were plotted together into a line graph. Additionally, to identify the best clustering for nodose ganglia neurones, the following steps were performed: 5) the silhouette widths calculated and repeated from step 2) were aggregated by clusters and averaged to get the per-cluster ASW; 6) the per-cluster ASW of clusters with proportion of cells expressing *Phox2b* > 15% were recorded; 7) the *Phox2b* positive cluster ASWs were aggregated and averaged based on corresponding PCs and resolutions, and plotted as a line graph. Based on the overall and *Phox2b* positive clusters’ ASWs, the clusters generated using 50 PCs and resolution = 1.4 was regarded as the optimal model with a total of 53 clusters. (Figure S1B, C).

### Defining cell types

The gene markers for neuronal cells were *Phox2b* for nodose ganglia neurons (NGN) and *Prdm12* for jugular ganglia neurons (JGN)^14^. For non-neuronal cells, *Emcn* was used for endothelial cells (EC)^14^, *Acta2* and *Ebf2* for Fibroblasts (FB)^57^, *Ptprc* for hematopoietic cells (HC)^58^, *Mpz* for myelinated glial cells (MGC)^59^, and *Apoe*, *Dbi*, and *Fabp7* for satellite glial cells (SGC)^14^. Additionally, a cluster was regarded as glial cells (GC) if the marker genes of MGC and SGC were both highly expressed (Table S11). The expression of each marker gene was visualized, and a histogram of expression levels was used to characterise cell clusters.

### Differential gene expression analysis

To identify the marker genes for each cluster, the default setting of FindAllMarkers function from Seurat (Version 4.3.1)^69^ was used. The DEGs between two clusters of cells were defined by the Wilcoxon Rank Sum test. Specificity score was calculated as in Steuernagel et al^34^ to identify relevant markers for cluster annotation: the DEGs with the highest per cluster were used as cluster annotations. Genes starting with *Gm*-, ending with -*Rik*, or genes used by previous clusters were excluded as per^34^.

Differentially expressed genes in cells or nuclei from *ad libitum* fed or overnight fasted, or from left or right nodose ganglia, were identified using the FindMarkers function of Seurat (Version 4.3.1)^69^ with the Wilcoxon Rank Sum test. The number of statistically significantly upregulated (P < 0.05; Average log_2_Foldchange > 0) and downregulated (Average log_2_Foldchange < 0) DEGs from each cluster were plotted.

### Downstream pathway analysis

The ingenuity pathway analysis (IPA, QIAGEN, Germany) was used to identify pathways that were upregulated or downregulated under fed and fasted conditions, or what pathways were differentially regulated in the left or right nodose ganglia. The DEG lists used for pathway analysis were from the two NGN clusters with highest number of statistically significantly DEGs (P < 0.05) either upregulated (NGN6, NGN9) or downregulated (NGN2, NGN11) in fasting. Similarly, when comparing left to right nodose ganglia, the DEG lists were from the three NGN clusters with the highest number of statistically significantly DEGs (P < 0.05) enriched in left nodose ganglia (NGN10, NGN17, NGN19), and the cluster with the highest number of DEGs (P < 0.05) enriched in right nodose ganglia (NGN5). The pathways with top 10 -log(B-H *p* value) from each cluster were grouped and plotted into a heatmap.

### Neurotransmitter/Neuropeptide assignment

Neuropeptides and neurotransmitters were assigned to individual neuronal clusters based on a set of criteria similar to that described in Langlieb et al^73^. For neurotransmitters, neuronal clusters were assigned neurotransmitter identity based on the percentage of cells in a cluster expressing marker genes required for the synthesis/transport of a neurotransmitter. In this instance, a threshold of 30% was used. If a cluster was not assigned any neurotransmitter identity, the expression of neurotransmitter related genes was inspected, and a neurotransmitter was manually assigned. Table S12 indicates the genes used to assign neurotransmitter identity to each cluster. For neuropeptides, expression was defined if over 30% of cells in a cluster had at least 1 transcript of the gene. Some neuropeptides were found to be highly expressed in all clusters, and so higher cut offs were chosen for these genes after manually inspecting a histogram of gene expression across all clusters. The following cut offs were used for high expressing genes: *Cartpt* > 60%; *Adcyap1*, *Calca*, *Calcb*, *Tac1*, *Bdnf* > 50%; *Gal* > 40%.

### Assigning neuronal properties

The neuronal clusters were further characterised into different subtypes based on the expression level of sodium channels, mechanosensor and nocisensors, and the marker genes of neuronal fibres and neuronal projection to peripheral organs (Table S14)^14,15,30^.

### Cell-cell communication

CellChat (2.1.2)^42^ was used to investigate the enriched signalling pathways between mouse nodose ganglia and hindbrain neurons. We extracted neuronal clusters from Dowsett et al^43^ and merged these with neuronal clusters from NodoMap to create a CellChat object with nodose, jugular and hindbrain neurons. We focused on ‘Non-protein Signaling’ and ‘Secreted Signaling’ pathways to look at neurotransmitter and neuropeptide interactions between nodose/jugular neurons and hindbrain neurons. Enriched pathways were subsetted to only include pathways that were signalling in the biologically appropriate direction (from nodose/jugular to hindbrain), and were split into classifications based the number of clusters each pathway was signalling from/to (many:many, many:few, few:many, few:few). For each enriched pathway, if the number of clusters it was signalling from or to was greater than 25% of the clusters from that region then it was classified as ‘many’. To focus on enriched pathways likely to be reaching regions within the dorsal vagal complex (DVC), expression of DVC-specific genes were inspected clusters from the hindbrain snRNASeq dataset to highlight populations that are most likely from the area postrema, nucleus of the solitary tract, and dorsal motor nucleus of the vagus. We looked at expression of *Phox2b, Glp1r, Gcg, Dbh, Prlh, Gfral* and identified HB_NE_Ddc/Sctr, HB_NE_Tbx20/Prph, HB_NE_Gcg/Prlr, HB_NE_Ddc/Dbh, HB_NE_Gal/Dkk2 and HB_NE_Tfap2b/Olfr78 as DVC specific clusters, and so focused on ligand-receptor pairs that involved these populations.

### Spatial Transcriptomics pre-processing

After demultiplexing, STAR (version 2.7.5)^74^ was used to align the spatial RNA sequencing data to the reference transcriptome GRCm38 (mm10). Following Curio Seeker pipeline (version_2.0.0)^75^, the detected cell-associated barcodes, features, and the feature counts of every barcode were generated and saved as rds files for downstream analysis. Each dataset was filtered to keep the spot with more than 100 transcripts and 190 genes.

### Integration of the spatial transcriptomics data of mouse left and right nodose ganglia

The integration of spatial transcriptomics datasets followed the same Seurat integration pipeline as the mouse nodose ganglia sc/snRNASeq data using sctransform (Version 0.3.4)^70^. The integrated nodose ganglia spatial transcriptomic data had 24,576 features/genes and 9,683 barcodes/spots.

### Integration of snRNASeq and spatial transcriptomics data

To visualise the location of different cell types across the nodose ganglia, the Robust Cell Type Decomposition (RCTD)^37^ pipeline (spacexr, version 2.2.1) was followed. RCTD is a computational method that uses a sc/snRNASeq data as a reference to characterise the cell types in a ST dataset. From the integrated mouse nodose ganglia sc/snRNASeq dataset, the untransformed count matrix, the cell cluster information (PC = 50, resolution = 1.4), and the total number of counts/transcripts of each cell were extracted as the reference. In the integrated ST data, the spatial coordinates list, the untransformed count matrix, and the total number of counts of each pixel were used as the query. With the create.RCTD and run.RCTD (doublet mode) functions from spacexr (version 2.2.1)^37^, the cell cluster information from the reference was used to annotate every spot of the query.

### Neighbourhood analysis to identify spatial patterns in cell distribution

To investigate the structure of the ganglia, a neighbourhood analysis was performed to identify which types of cells commonly appeared together. Firstly, an artificial grid was created to bin the neighbouring spots together. The number of spots in each bin ranged between 0 and 20. Next, a matrix containing the bin ID and each spots cell type identification (from the RCTD output) was created: for spots identified as singlets only the first cell type was counted, for spots identified as doublet_certain and doublet_uncertain, both cell types were counted, and for spots identified as reject, no cell types were counted. Principal component analysis and clustering were applied to the count matrix to identify the types of neighbourhoods.

## KEY RESOURCES TABLE

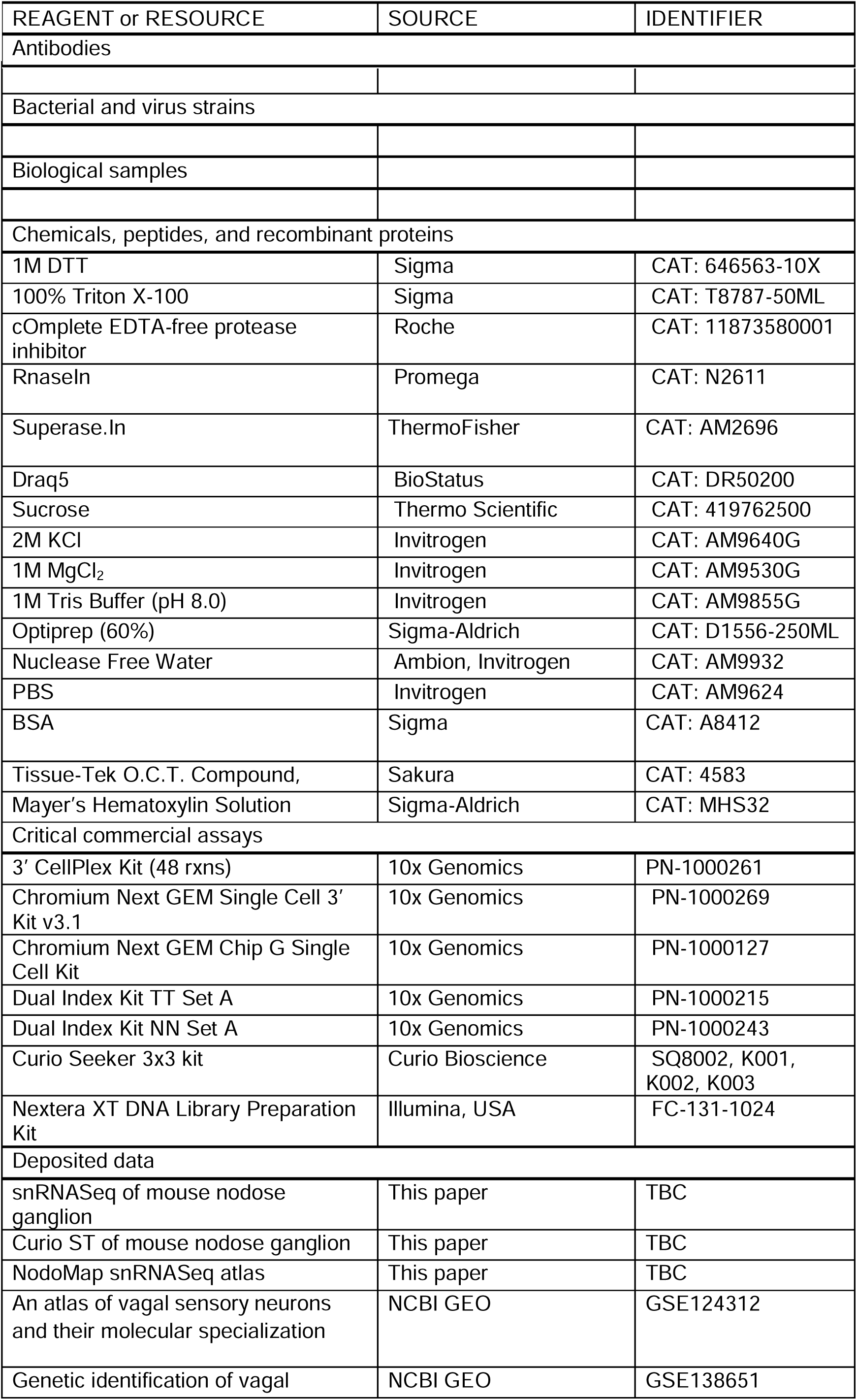
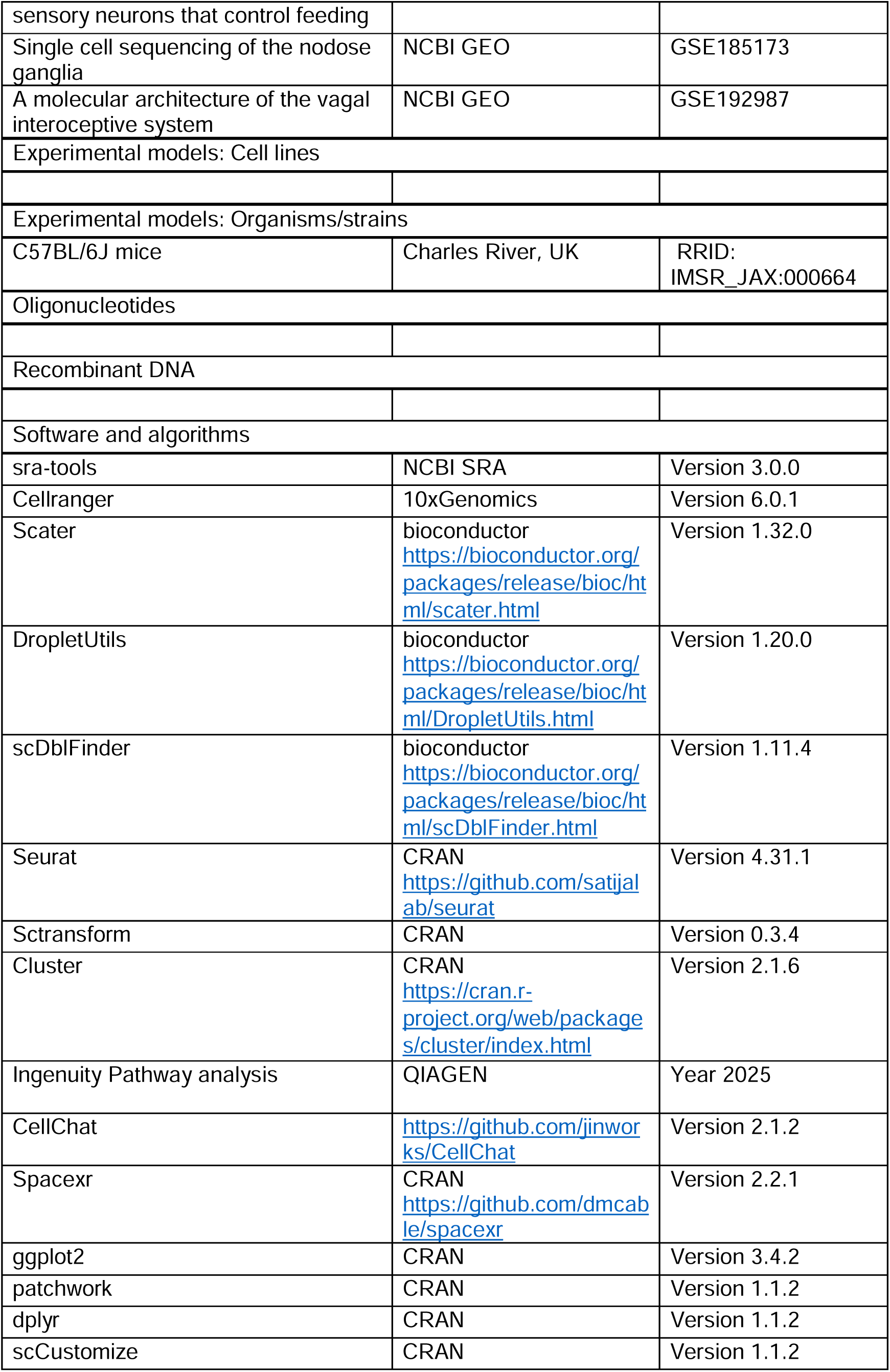
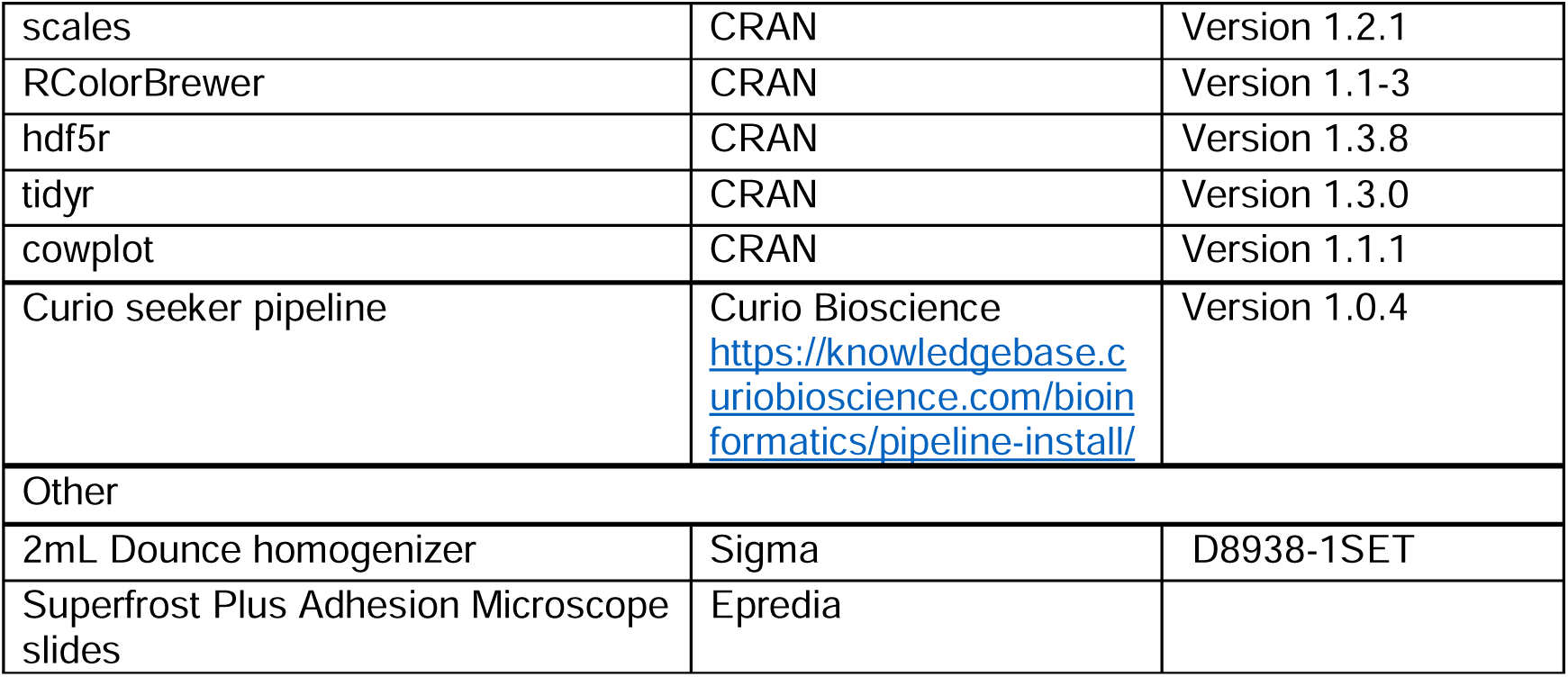

## 5. Data availability

Raw sequencing files generated in this manuscript are available on GEO at GSE296454.

## 6. Code availability

All code for the pre-processing, integration and plotting of data is available on GitHub at: https://github.com/sc2470/NodoMap

---

**Supplementary Figure 1:**
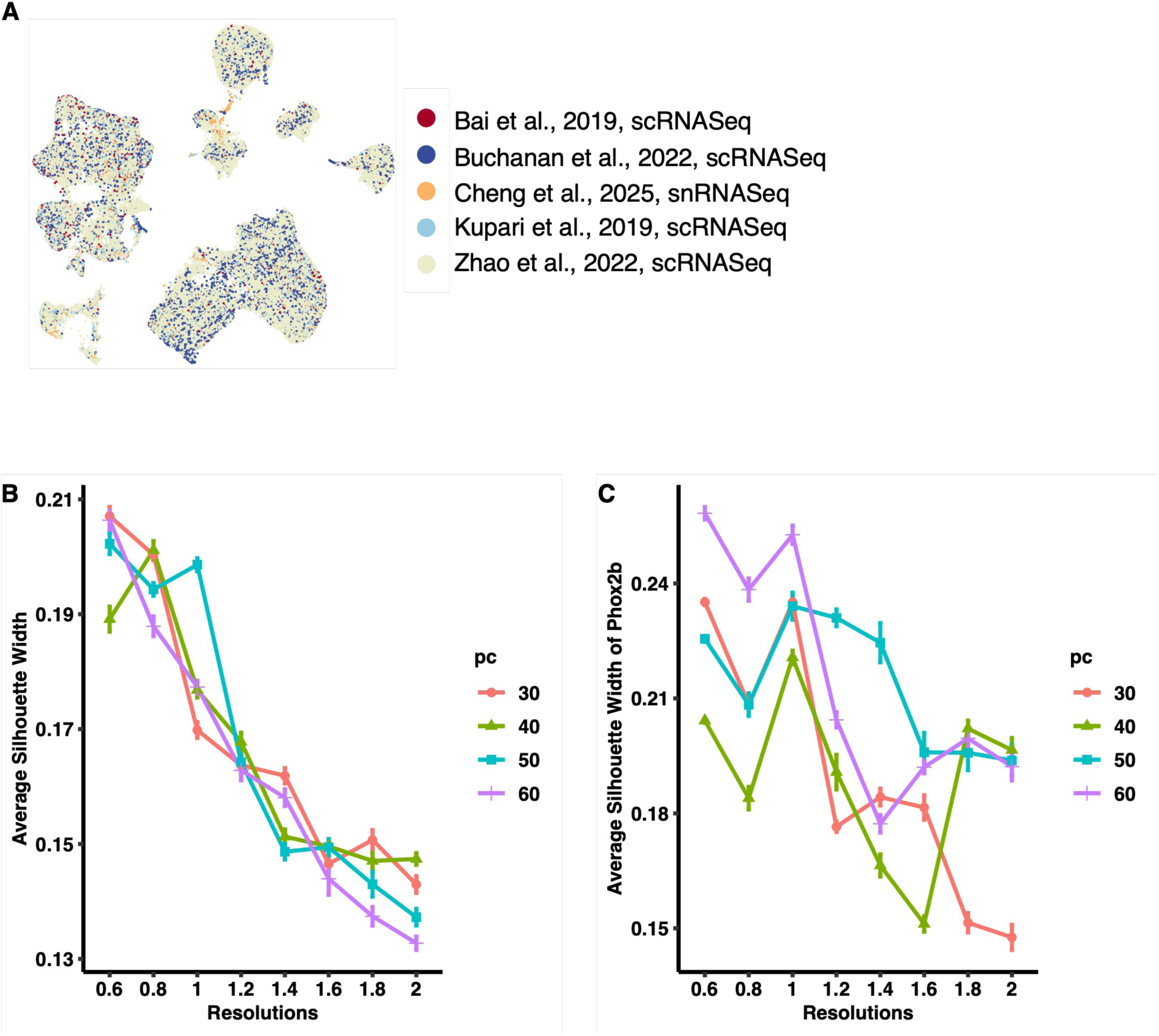
Dataset integration quality control. **(A)** UMAP plot coloured by dataset. There were 2,079 from Bai et al^15^, 17,892 from Buchanan et al^28^, 1,955 from Cheng et al, 4,310 from Kupari et al^14^ and 82,246 from Zhao et al^30^ single cells/nuclei used for downstream analysis. **(B-C)** Line plots of averaged average silhouette width score of all clusters **(B)** and *Phox2b* expressed clusters **(C)** from principal components (PCs) 30 to 60 (increment = 10) and resolutions 0.6 to 2 (increment = 0.2). Together with log-normalised expression of *Phox2b* (NGN marker) and *Prdm12* (JGN marker), single cell atlas defined under PC = 50 and resolution = 1.4 was regarded as the best clustering model.

**Supplementary Figure 2:**
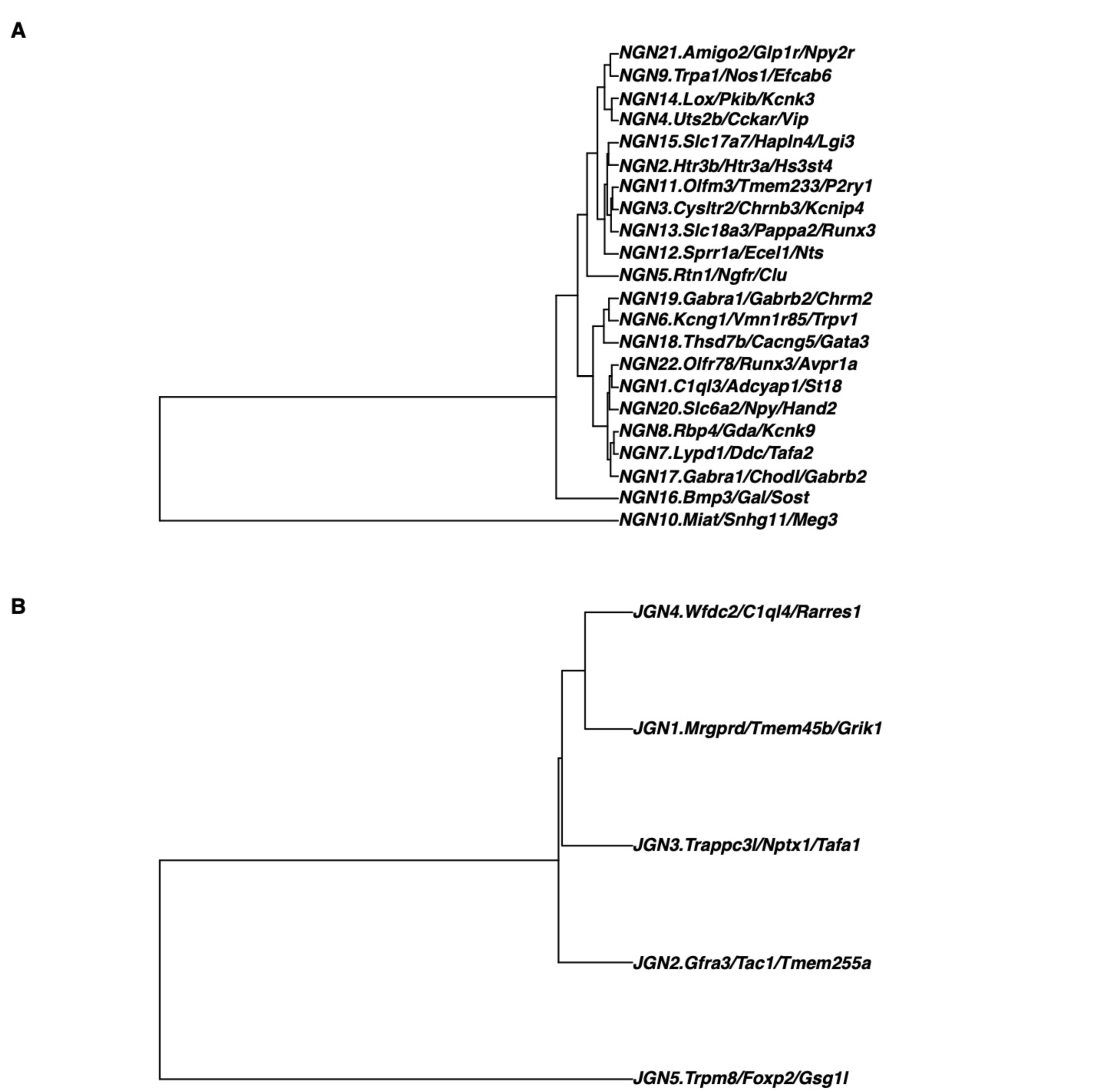
Dendrograms of the neuronal clusters. **(A)** The phylogenetic tree plot of the 22 NGN clusters. **(B)** The phylogenetic tree plot of the 5 JGN clusters.

**Supplementary Figure 3:**
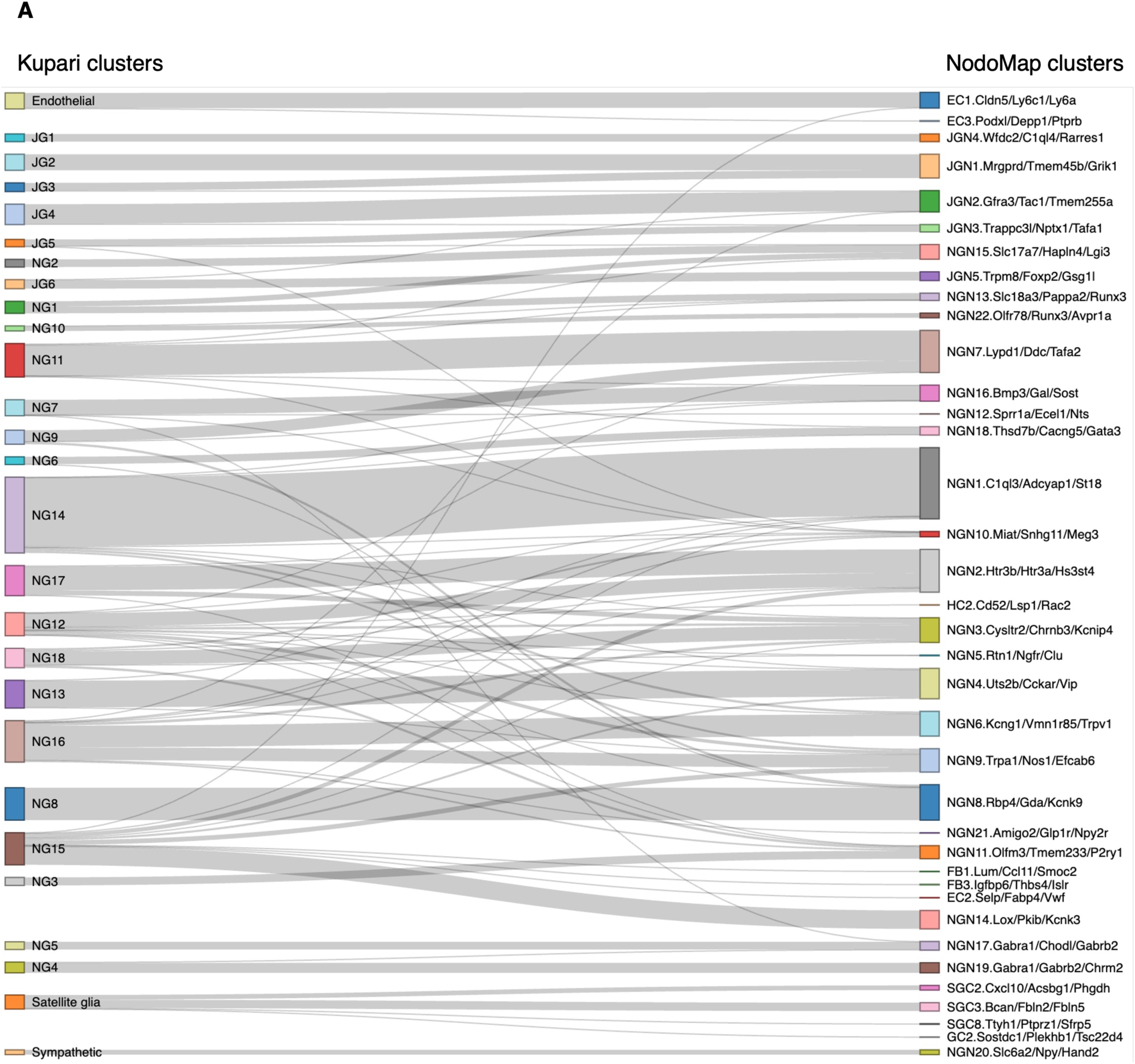
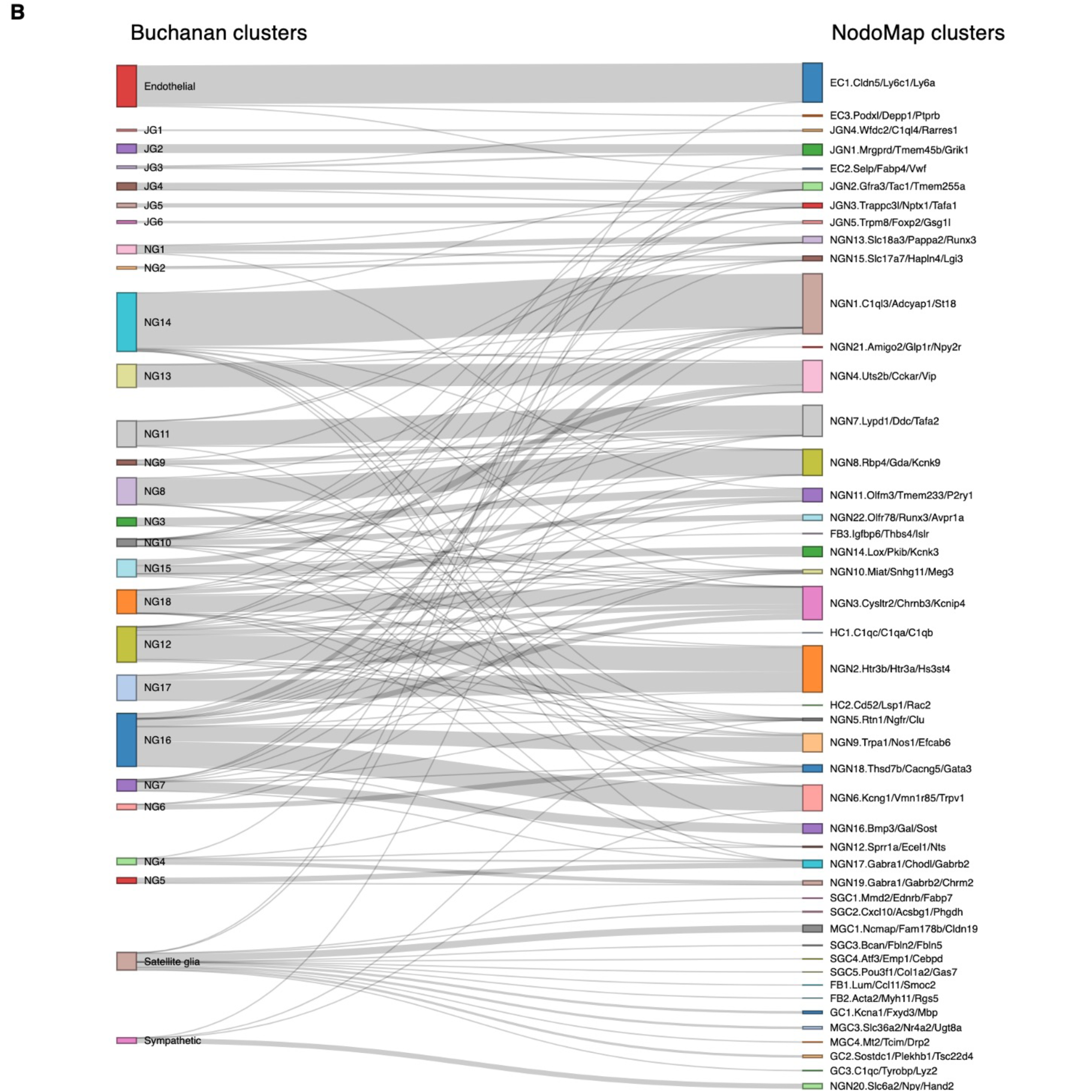
**(A)** Sankey plot highlighting the alignment of the cluster annotation from Kupari et al^14^ compared with the annotation based on all five datasets (NodoMap clusters). **(B)** Sankey plot aligning the cluster annotation from Buchanan et al^28^ and NodoMap.

**Supplementary Figure 4:**
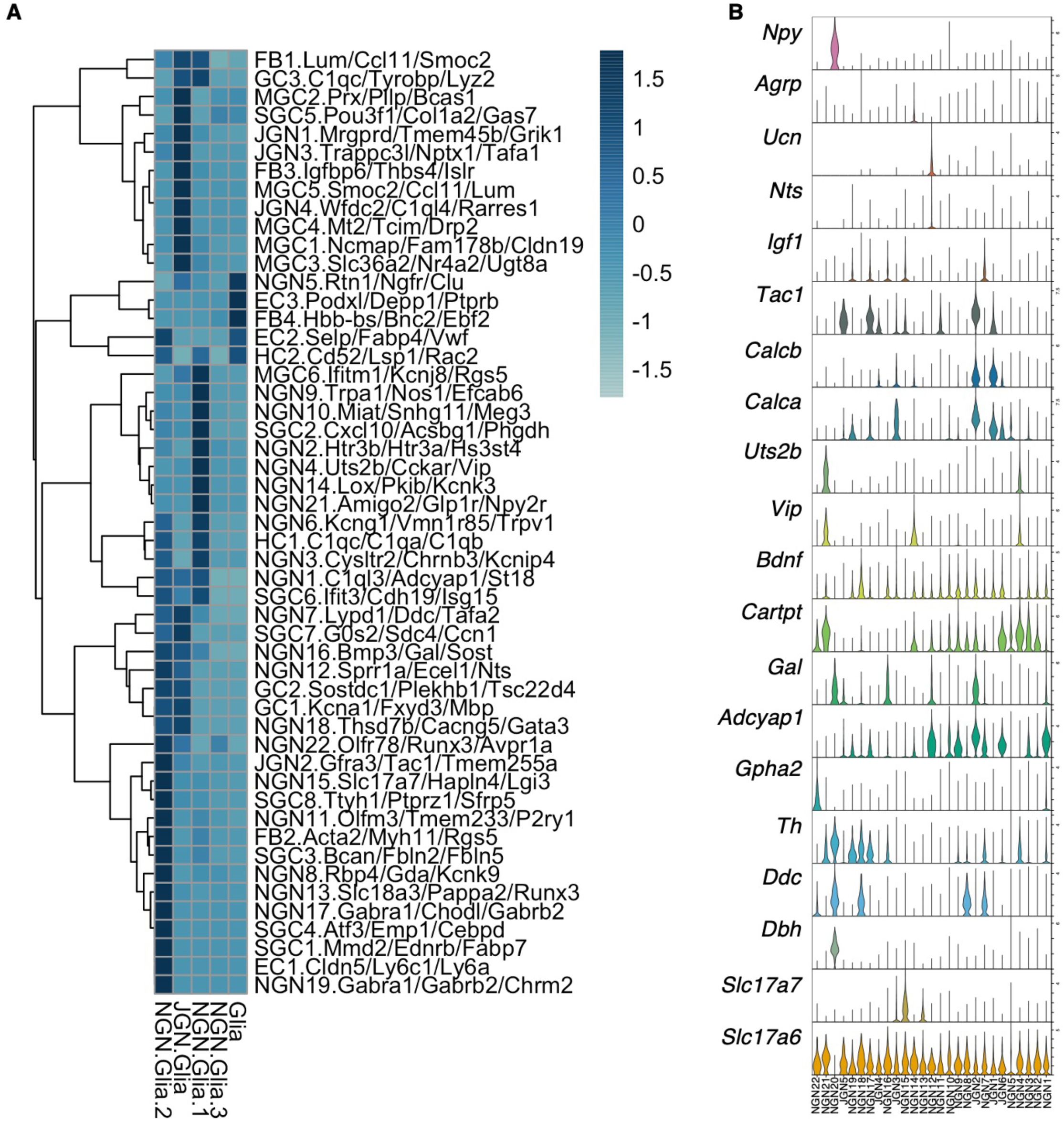
**(A)** Neighbourhood analysis was performed on the RCTD cell assignments in the spatial transcriptomics data. This formed a total of five neighbourhoods which were then labelled based on the co-occurrence of different cell types. The heatmap displays the scaled appearance of assigned cell types within each bin in the spatial transcriptomics dataset. Rows are clustered. **(B)** Violin plot showing expression of neuropeptide- and neurotransmitter-associated genes in nodose and jugular neuronal clusters.

**Supplementary Figure 5:**
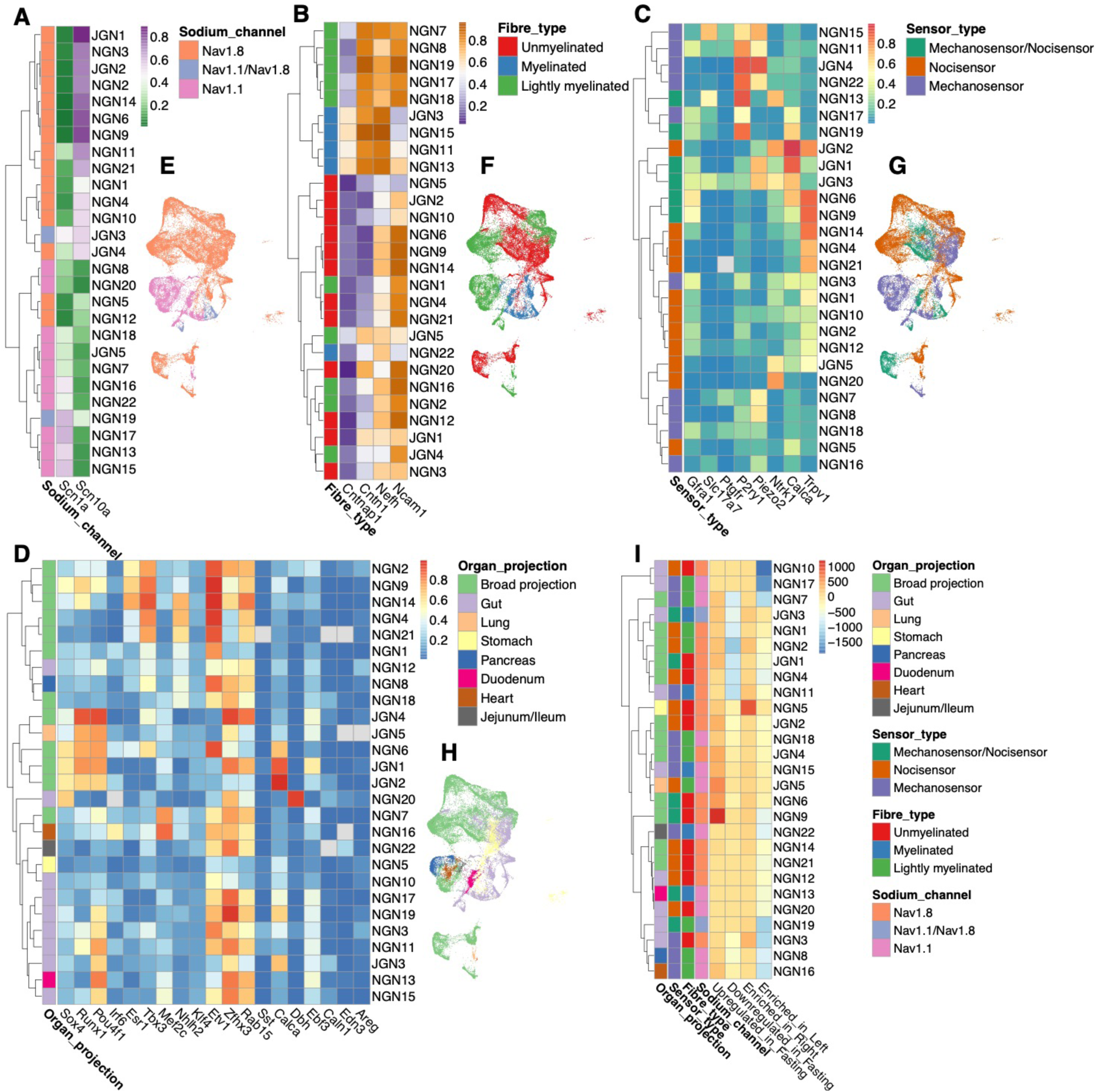
Neuronal cluster classifications. **(A-D)** The clustered heatmaps showing the proportion of cells expressing the marker genes within the corresponding neuronal clusters. The neuronal clusters were clustered by the Boolean values. By comparing the expression level of marker genes, the neuronal clusters were further characterised based on their **(A)** sodium channels (Nav1.1 (*Scn1a*) or Nav1.8 (*Scn10a*)); **(B)** fibre types (myelinated or unmyelinated (*Cntnap1*, *Cntn1, Nefh, Ncam1*)); **(C)** sensor types (mechanosensor (*Gfra1, Slc17a7, Ptgfr, P2ry1, Piezo2*) or nocisensor (*Ntrk1, Calca, Trpv1*)); **(D)** organ projections (lung (*Sox4, Runx1, Pou4f1*), heart (*Irf6, Esr1, Tbx3, Mef2c*), pancreas (*Nhlh2, Klf4*), gut (*Etv1, Zfhx3, Rab15*), stomach (*Sst, Calca*), duodenum (*Dbh, Ebf3*), jejunum/ileum (*Caln1, Edn3*), broad projection (marker genes from two or more organs). **(E-H)** The dimplots of NodoMap neuronal clusters, coloured based on the classification of sodium channels **(E)**, fibre types **(F)**, sensor types **(G)**, and the peripheral organ projections **(H)**. **(I)** The heatmap of 4 neuronal annotations and the number of upregulated/downregulated DEGs in fasting or enriched DEGs in left or right nodose ganglia.

**Supplementary Figure 6:**
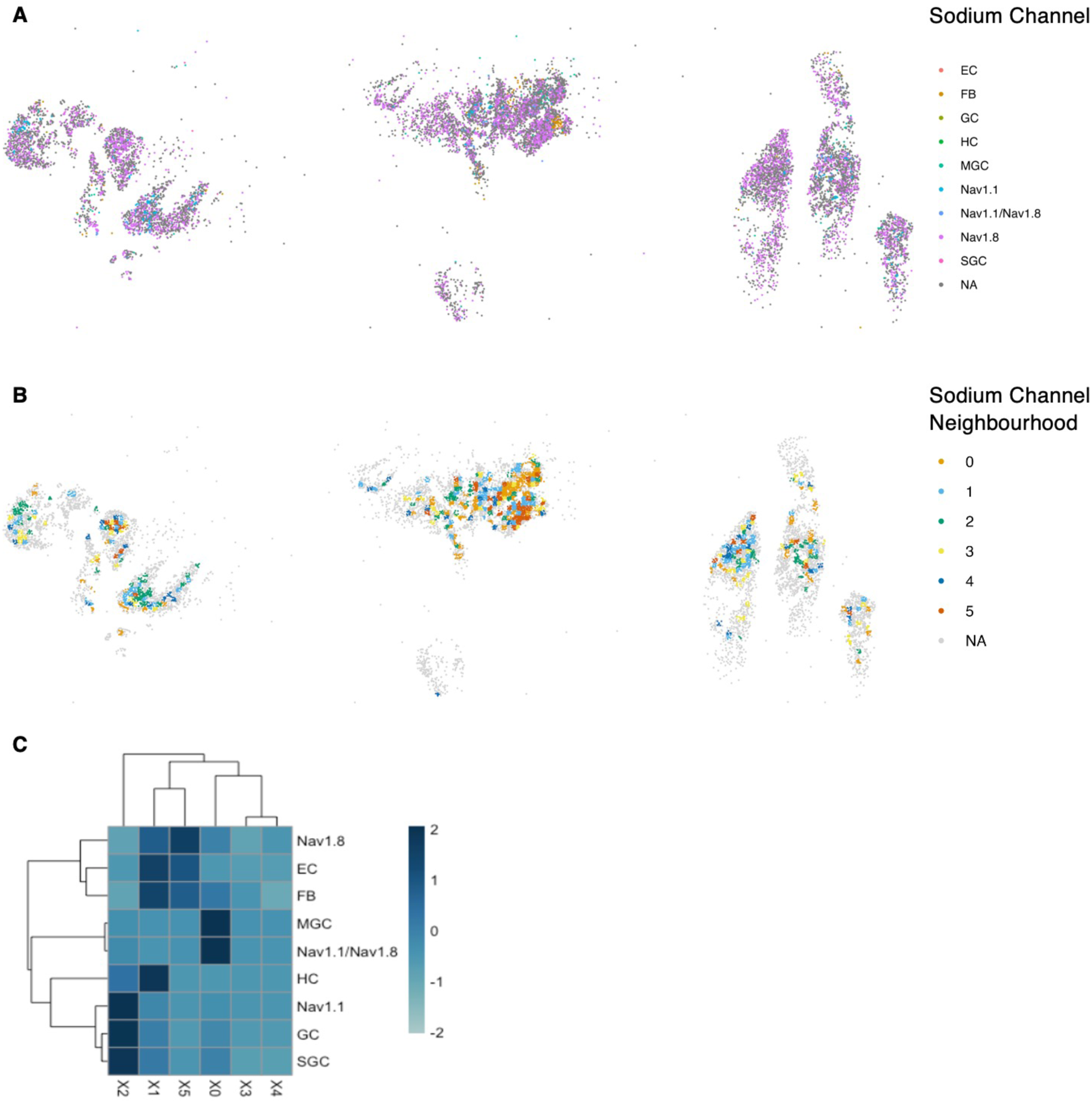
**(A)** RCTD annotation of spatial transcriptomics spots, coloured by sodium channel type. The left and central tiles each were mounted with a section from three left nodose ganglia. The right tile was a section from three right nodose ganglia. **(B)** Neighbourhood analysis of sodium channels and non-neurons identifies 6 neighbourhoods. **(C)** Heatmap showing scaled prevalence of each cell type in the neighbourhoods

**Supplementary Figure 7:**
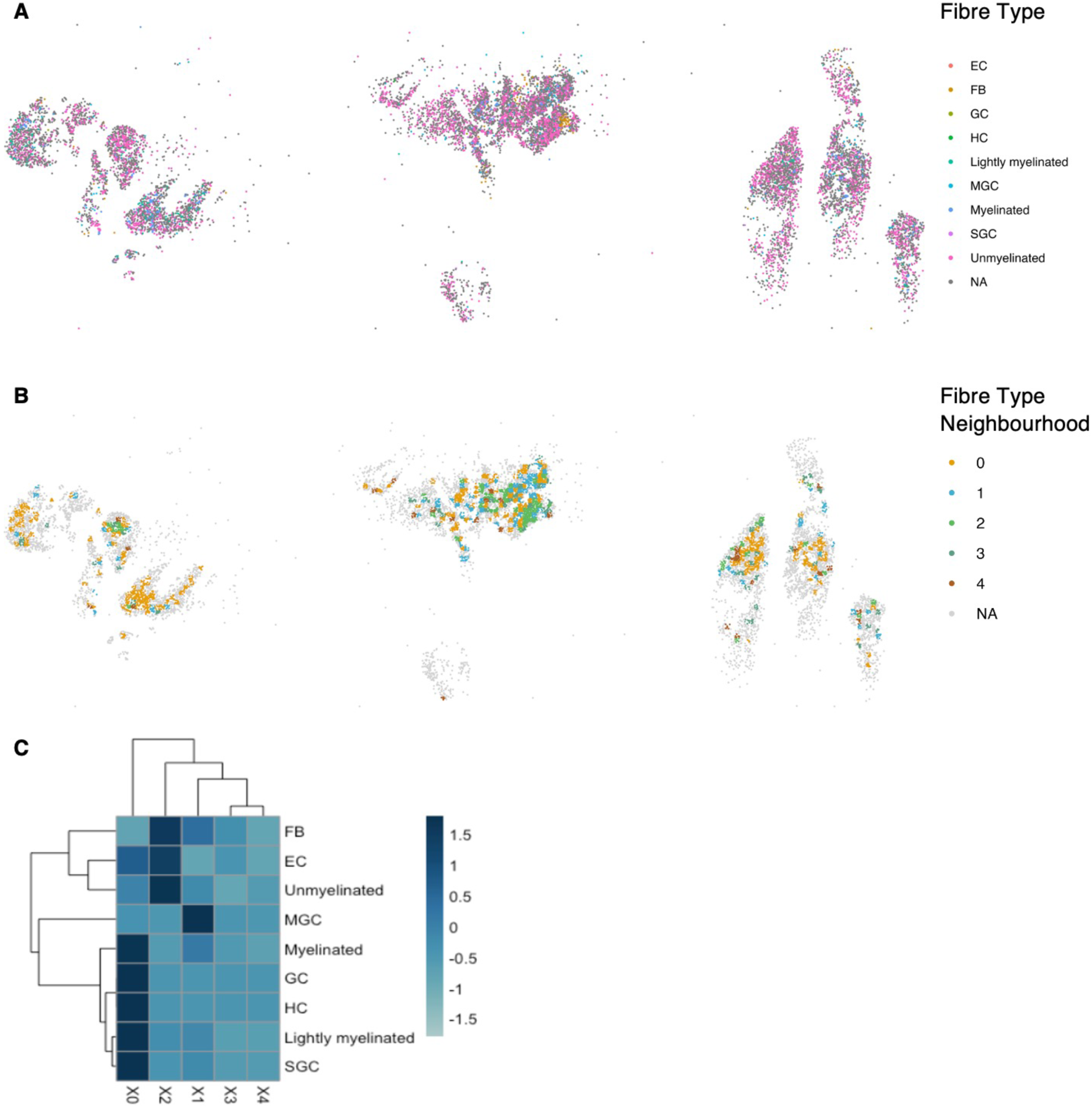
**(A)** RCTD annotation of spatial transcriptomics spots, coloured by fibre type. The left and central tiles each were mounted with a section from three left nodose ganglia. The right tile was a section from three right nodose ganglia. **(B)** Neighbourhood analysis of fibre types and non-neurons identifies five neighbourhoods. **(C)** Heatmap showing scaled prevalence of each cell type in the neighbourhoods

**Supplementary Figure 8:**
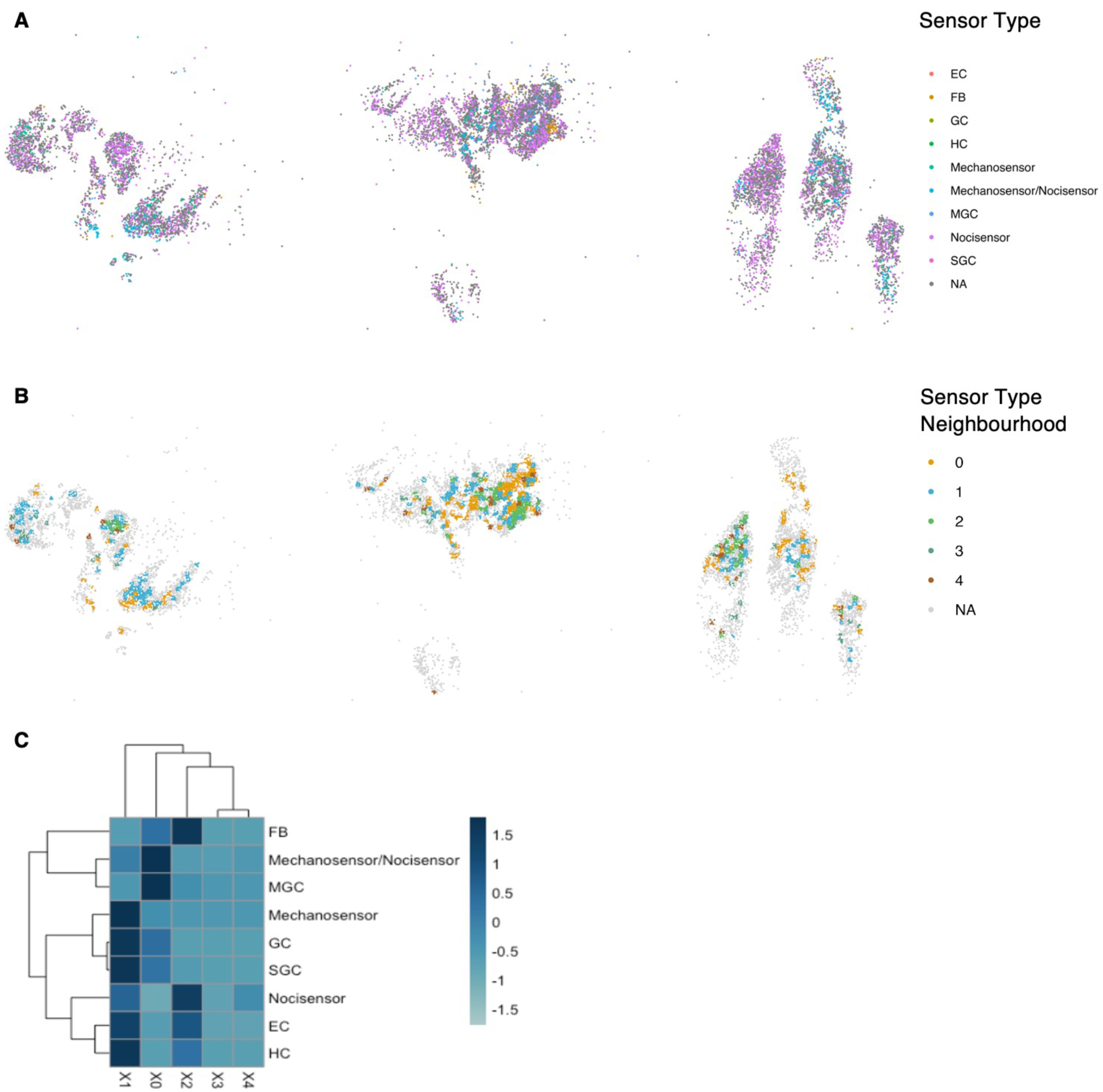
**(A)** RCTD annotation of spatial transcriptomics spots, coloured by sensor type. The left and central tiles each were mounted with a section from three left nodose ganglia. The right tile was a section from three right nodose ganglia. **(B)** Neighbourhood analysis of sensor types and non-neurons identifies five neighbourhoods. **(C)** Heatmap showing scaled prevalence of each cell type in the neighbourhoods

**Supplementary Figure 9:**
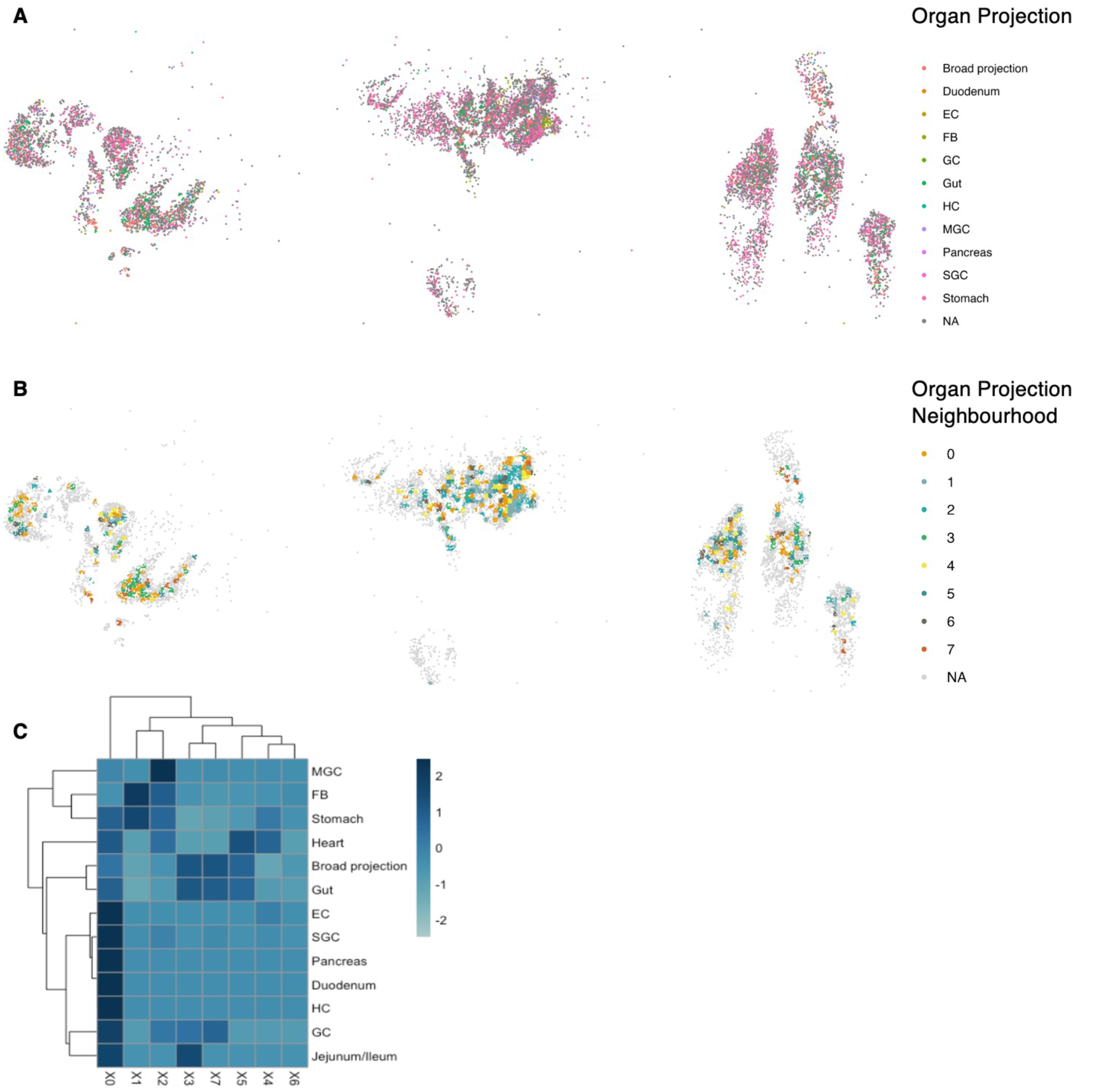
**(A)** RCTD annotation of spatial transcriptomics spots, coloured by organ projection annotation. The left and central tiles each were mounted with a section from three left nodose ganglia. The right tile was a section from three right nodose ganglia. **(B)** Neighbourhood analysis of organ projection annotation and non-neurons identifies eight neighbourhoods. **(C)** Heatmap showing scaled prevalence of each cell type in the neighbourhoods

**Supplementary Figure 10:**
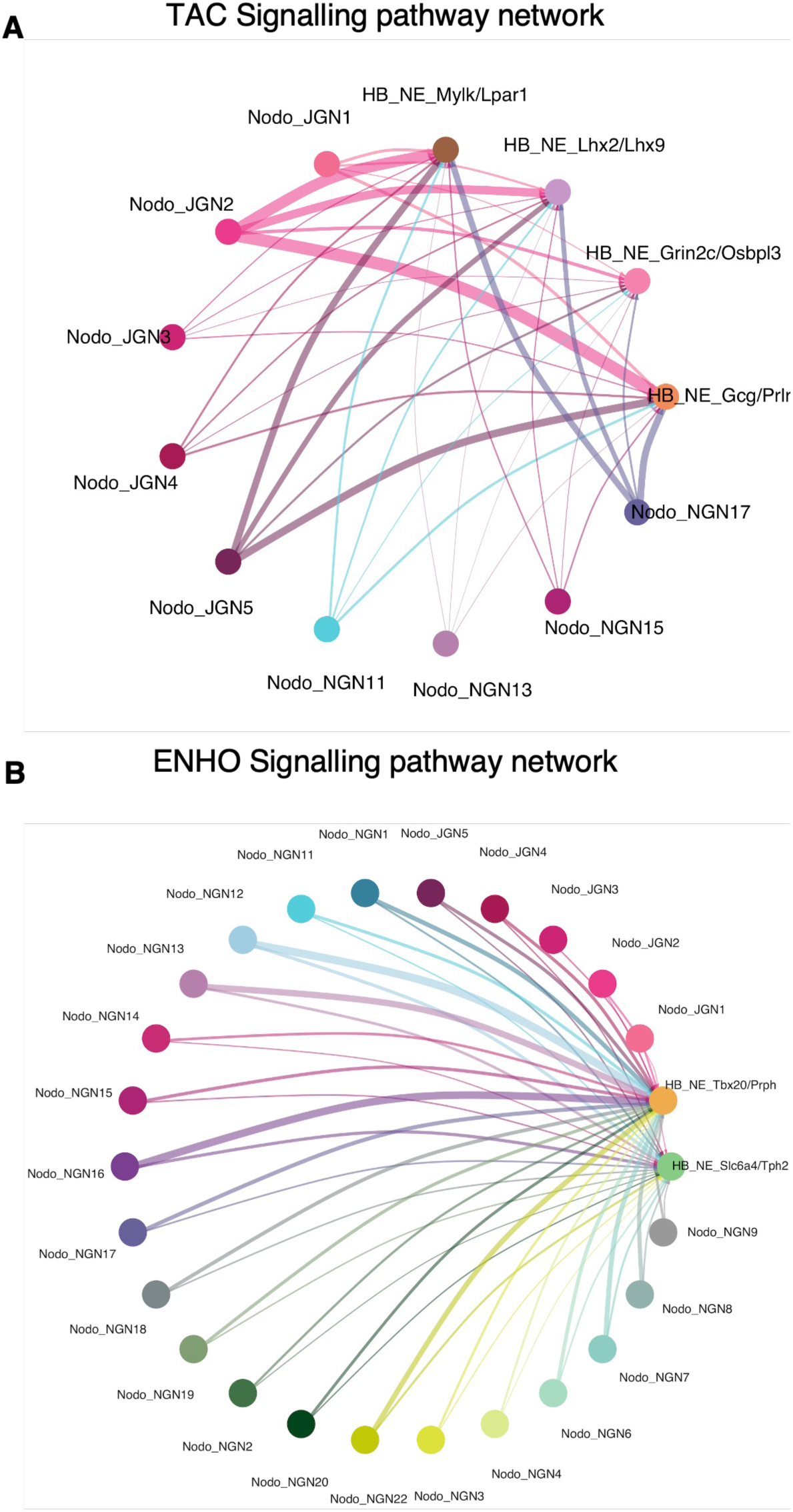
CellChat mapping showing ligand-receptor interactions between nodose neuronal clusters and hindbrain neuronal clusters for TAC signalling (top) and ENHO signalling (bottom). The thickness of the line is an indication of the estimated strength of the signalling.

